# Mapping HIV-1 RNA Structure, Homodimers, Long-Range Interactions and persistent domains by HiCapR

**DOI:** 10.1101/2024.09.03.610837

**Authors:** Yan Zhang, Jingwan Han, Dejian Xie, Wenlong Shen, Ping Li, Jian You Lau, Jingyun Li, Lin Li, Grzegorz Kudla, Zhihu Zhao

## Abstract

Human Immunodeficiency Virus (HIV) persists as a leading global health issue. A significant gap in knowledge exists in our understanding of long-range interactions within the HIV-1 RNA genome. To bridge this gap, we introduce HiCapR, incorporating a psoralen crosslinking RNA proximity ligation and post-library hybridization for capturing HIV RNA-RNA interactions.

Leveraging HiCapR, we confirm the presence of stem structures in the key regions, such as the 5’-untranslated region(5’-UTR) and Rev Response Element (RRE) stems, and dimer sites in 5’-UTR region, which is responsible for HIV packaging. Importantly, we reveal multiple previously unknown homodimers along the HIV genome, which may have important implications for viral RNA splicing and packaging processes. Also, we uncover a wealth of unprecedented long-range interactions, particularly within the 5’-UTR of infected cells.

Intriguingly, our findings indicate a pronounced reduction in long-range RNA-RNA interactions, signifying a transition from a state of abundant interactions, hence a relative loose state within infected cells to a condensed structure within virions. Concurrently, we have demonstrated the presence of stable genomic domains within virions that are instrumental in the dimerization process. These domains are preserved throughout the packaging process.

Our findings shed light on the functional significance of RNA organization, including stable and persistent genomic domains, homodimerization, and long-range RNA-RNA interactions, in the splicing, packaging as well as assembly of HIV.

**Highlights:** HiCapR is a new proximity ligation method for mapping RNA structures and homodimers in the HIV genome with sufficient reliability and efficiency.

Multiple homodimers were discovered along the genome, with potential implications for splicing and packaging processes.

Long-range RNA-RNA interactions are abundant in infected cells but significantly reduced in virions.

Stable genomic domains encluding homodimer sites are persistent in virions and are involved in dimerization.

## Main

Human Immunodeficiency Virus (HIV) remains a significant global health concern despite advancements in antiretroviral therapy even with the newest report of total seven AIDS patients been completely “cued” ^1 2,3^. Research on HIV-1 RNA structure and function has intensified, focusing on assembly, release, and maturation processes, using NMR^4^, cryo-EM^5^ and chemical probing^6-8^. However, our knowledge of the global architecture and the long-range interactions within the HIV-1 RNA genome remains limited, primarily due to the complexity and dynamic nature of these structures.

The HIV genome, comprising two copies of positive-sense RNA, has been extensively studied to understand its regulatory elements, such as the 5’-UTR, crucial for dimerization and selective packaging into viral particles^9^. Additionally, interactions involving the nucleocapsid protein (NC) and the dimerization signal (Ψ) play pivotal roles in the assembly and maturation of HIV virions^10^. Furthermore, the RRE in the HIV genome, responsible for nucleocytoplasmic transport of viral RNA, exhibits structural variability impacting virus replication rates^11,12^. While current techniques like SHAPE began to provide insights into local RNA structures^7,8,13^, studying the intricate long-range RNA interactions and homodimers within the HIV genome under physiological conditions remains a significant challenge, primarily due to the limitations of these methods in capturing the full spectrum of RNA interactions and the dynamic nature of the viral genome.

Proximity ligation-based methods have been shown to be particularly useful for studying long-range RNA interactions, as they can overcome the RNA length limitations of traditional techniques. These methods have been applied to the study of various viruses, including influenza^14,15^, Zika^16-18^, and SARS-CoV-2^19-22^, and have led to the identification of numerous RNA structures that are closely related to the virus lifecycle. We and others have recently shown that RNA proximity ligation data also contains information about RNA homodimerization events^23,24^. Nevertheless, as of now, there are no proximity ligation studies of long-range interactions or dimeric interactions of the HIV RNA genome. Probably due to high requirements of starting material, making it very difficult for low abundant HIV sample treatment.

In this study, we modified our previous protocol^22^ by integrating a hybridization-based capture of HIV-1 sequence after library construction, resulting in a method we call High throughput Capture of RNA interactions (HiCapR). This advancement enabled the comprehensive capture and analysis of the complete HIV RNA genome. By applying HiCapR to both infected cells and virions, we uncover a distinct genomic compression pattern in virions, highlighting the critical role of global genome folding in HIV packaging and assembly. Our findings not only reveal the presence of persistent genomic domains within virions that facilitate whole genome dimerization, even the transgenerational inheritance but also a significant reduction of long-range RNA-RNA interactions in virions compared to infected cells. These insights provide a new perspective on the functional implications of HIV RNA structure in splicing and packaging processes and may lead to the identification of new targets for antiviral intervention.

## results

### 1. Capturing in vivo HIV genome RNA structure in infected cells and in virion

To effectively capture and analyze full-length low abundant HIV RNA-RNA interactions in infected cells, we developed an improved version of the protocol, which we call High throughput Capture of RNA interactions (HiCapR). This method incorporates hybridization-based capture with next-generation sequencing (NGS) library construction, a principle well established in capture Hi-C methods ^25-28^ and in RNA proximity ligation ^17,19^. Applying this strategy, we compared the NL4-3 and GX2005002 strains ^29^,which are two distinct HIV-1 strains that exhibit different prevalence patterns in various geographical regions. The NL4-3 strain is a well-characterized laboratory strain that is widely used in HIV research and is representative of the HIV-1 subtype B, which is highly prevalent in Europe and the Americas. On the other hand, GX2005002 is a primary isolate of the CRF01_AE subtype, which is one of the most prevalent strains in Southeast Asia, particularly in China. The overall design of this study is presented in Figure 1A, with detailed methods provided in the methods section. Supplementary Figure S1 illustrates the bioinformatic pipeline used to detect RNA-RNA interactions and homodimers.Approximately 16 million raw reads were obtained per sequencing library, with over 95% of the reads containing HIV sequences, indicating efficient capture of HIV RNA transcripts. The alignment rate exceeded 90%, and we achieved an average depth of coverage of approximately 11,500X across the genome (Supplementary Table 1). The uniform distribution of reads post-capture (Supplementary Figures S2) demonstrates HiCapR’s effectiveness in capturing the entire HIV RNA genome with minimal bias. The overall duplication rate remained below 50%, with a lower rate in the ligation group, suggesting increased library complexity due to RNA proximity ligation.

**Figure 1.**
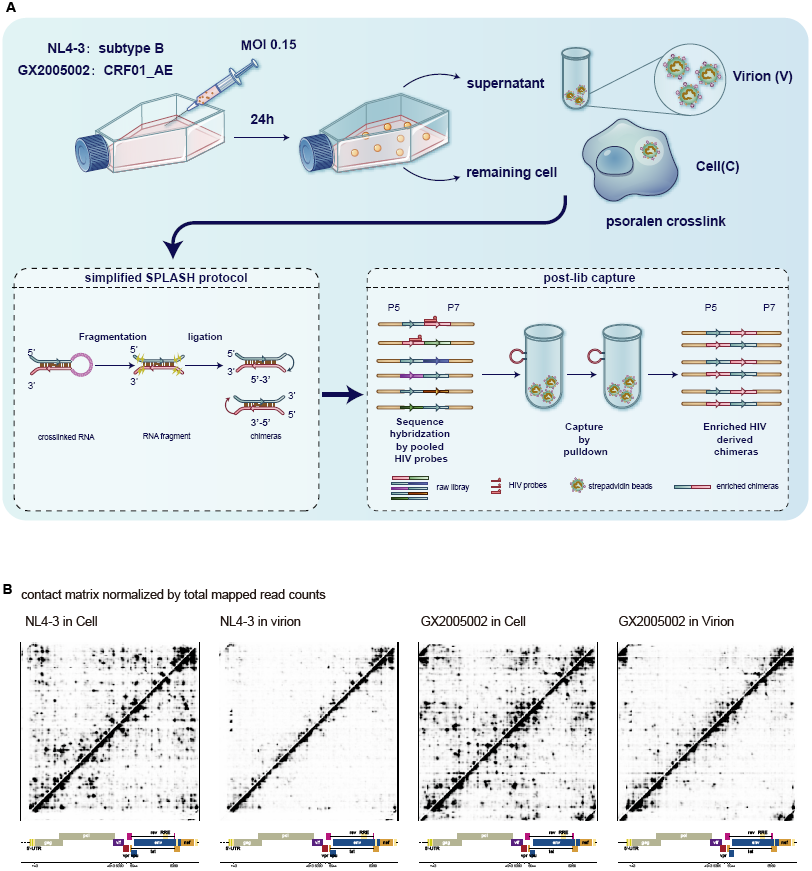
Experimental design of HiCapR and overall HIV-1 contact matrix. A) Experimental design of HiCapR for profiling RNA-RNA interactions and dynamics of the HIV genome. Based on the simplified SPLASH protocol, HiCapR incorporates a post-library probe-based hybridization and streptavidin pulldown method to enrich HIV RNA chimeras from the SPLASH library. The HiCapR method has been applied to two strains of HIV, NL4-3 and GX2005002, in both infected cells and their corresponding virus particles. B) Comprehensive contact matrix derived from infected cells and virions of both NL4-3 and GX2005002 HIV-1 strains in the HiCapR experimental groups. The heatmap displays the average count of chimeras per 1 million mapped reads, combining data from two biological replicates for each sample.

Chimeric reads were identified using the hyb technique as previously described^22^, with a database comprising human spliced mRNAs, noncoding RNAs^30^ and the HIV genome sequence as reference. The chimeric rate, a crucial indicator of ligation efficiency, was approximately 1% in non-ligation controls and around 9% in the ligation group, aligning with the widely reported range. The proportion of 5’-3’ and 3’-5’ chimeras was nearly equal in long-range interactions, while proximal interactions were enriched for 3’-5’ chimeras. Pearson correlation analysis revealed a high degree of correlation (r>0.99) between biological replicates, indicating high reproducibility. Additionally, a significant similarity was observed between virions in the supernatant and infected cells from the same viral strain (Supplementary Figure S3).

Using a similar approach as described in our previous paper ^22^, we generated a contact matrix for the global HIV-1 genome, revealing clear local and long-range interactions (Figure 1B and Supplementary Figure S4). In summary, HiCapR demonstrated high reproducibility, low bias, reliability, and efficiency in capturing HIV transcripts, making it a robust tool for analyzing the RNA structure of the HIV genome and potentially other similar RNA viruses.

### 2. Local RNA structure heterogeneity, dynamics and robustness in HIV 5’-UTR

We aimed to characterize specific local structures and their dynamics within the HIV genome. One of the extensively investigated structures is 5’-UTR, as these structures are closely associated with crucial life processes of HIV.

Chimeras analysis support the presence of key structural elements, including TAR, polyA, SL1, SL2, and SL3, as well as polyA-SL1 in the monomeric conformation in the HIV genome (Figure 2A and Supplementary Figure S5). Despite the pivotal role of the 5’-UTR in replication, packaging, transcription and translation, we noted that the 5’-UTR sequence of HIV is not highly conserved across different HIV strains. A comparison between the NL4-3 and GX2005002 strains revealed notable insertions and deletions in the U5 region, along with point mutations in the loop of the SL1 core region, which is critical for HIV dimerization (Figure 2B). These findings raise questions about the conservation of the structural integrity of this region. We therefore applied the comradesFold algorithm ^17^ to our HiCapR data to generate 1000 structure predictions of the 5’-UTR (extending 100 nt downstream of the AUG start codon) for each sample, and then performed MDS analysis after aligning the viral genome coordinates of both strains (Figure 2C). This analysis revealed that the reported dimer and monomer structures clustered together in NL4-3 but form distinct clusters in GX2005002(Figure 2C). Based on these folded structures, we calculated the base pairing probability for each base pair, which involves determining the number of folded structures supporting a specific base pair divided by the total number of structures. Visualizing this base pairing probability as a heatmap identifies the most stable base pairs in the 5’ UTR of HIV. We observed a consistent presence of key structural elements such as polyA, TAR, SL1, SL2, and SL3 in both NL4-3 and GX2005002 strains (Figure 2D), suggesting robustness in the overall structure despite sequence variations and alternative RNA conformations.

**Figure 2.**
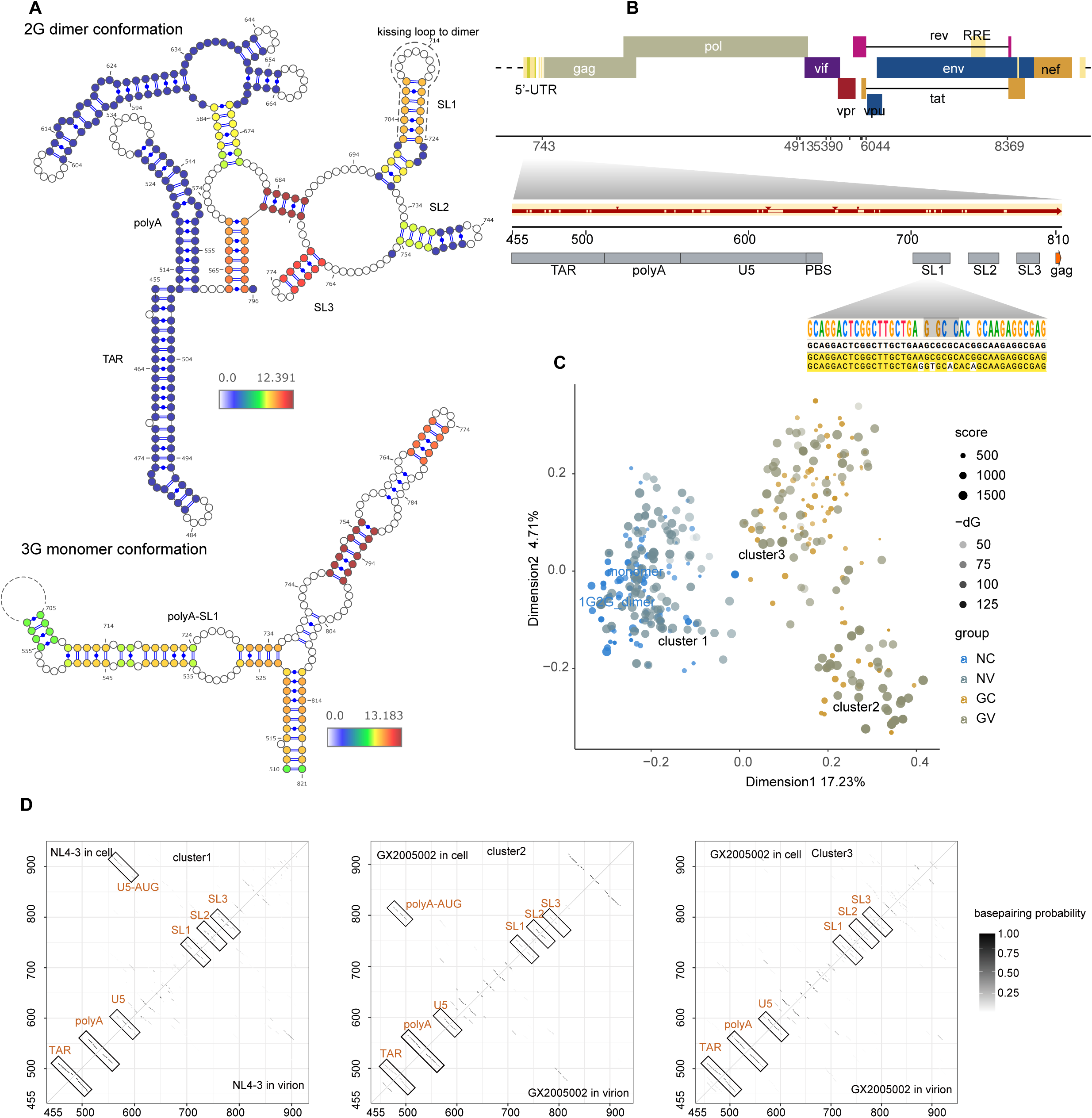
conformations of HIV-1 5’-UTR. A) Known conformation of HIV-1 5’-UTR supported by chimeras. Previous reported stems in 5’-UTR are supported by chimeras from HiCapR. Colors of the nucleotides indicate the log2 transformed base pairing scores. B) Representation of the one-dimensional structure of the HIV-1 5’-UTR, highlighting the conservation between the GX2005002 and NL4-3 strains in this region. The diagram includes rectangular boxes denoting the locations of key structural elements, with numerical coordinates referencing NCBI DNA genome coordinates. Dashed boxes indicate regions that are either absent or distinct in GX2005002 compared to NL4-3. Small triangle arrows indicate insertions in the GX2005002 5’-UTR. At the top, a seqlogo displays the consensus nucleotides in the SL1 region. C) Multidimensional scaling (MDS) plot clustering each of the 1,000 computationally predicted structures of the 5’-UTR in two strains under two conditions. D) Basepairing probability matrices were calculated from 1000 computationally predicted structures, with color indicating the percentage of structures supporting the specific basepair.

### 3. Novel homodimerization sites and their implications in RNA splicing and viral assembly

Proximity ligation techniques enable unambiguous detection of RNA homodimers from “overlapping” chimeric reads^23,24^ (Figure 3A). To ensure the accuracy of our results, we implemented a rigorous data filtering process to select chimeras formed exclusively through dimerization, minimizing background interference caused by RNA interactions or self-ligation (Supplementary Figure S1 and Methods). We quantified the total number of reads for dimeric chimeras, 3’-5’ chimeras, and 5’-3’ chimeras respectively in each sample (Supplementary Figure S6) and plotted contact matrices made from non-overlapping chimeras and dimeric chimeras (Supplementary Figure S7). The non-overlapping chimeras reveal short-range intramolecular RNA-RNA interactions, while dimeric chimeras capture homodimer formation in the HIV genome.

**Figure 3.**
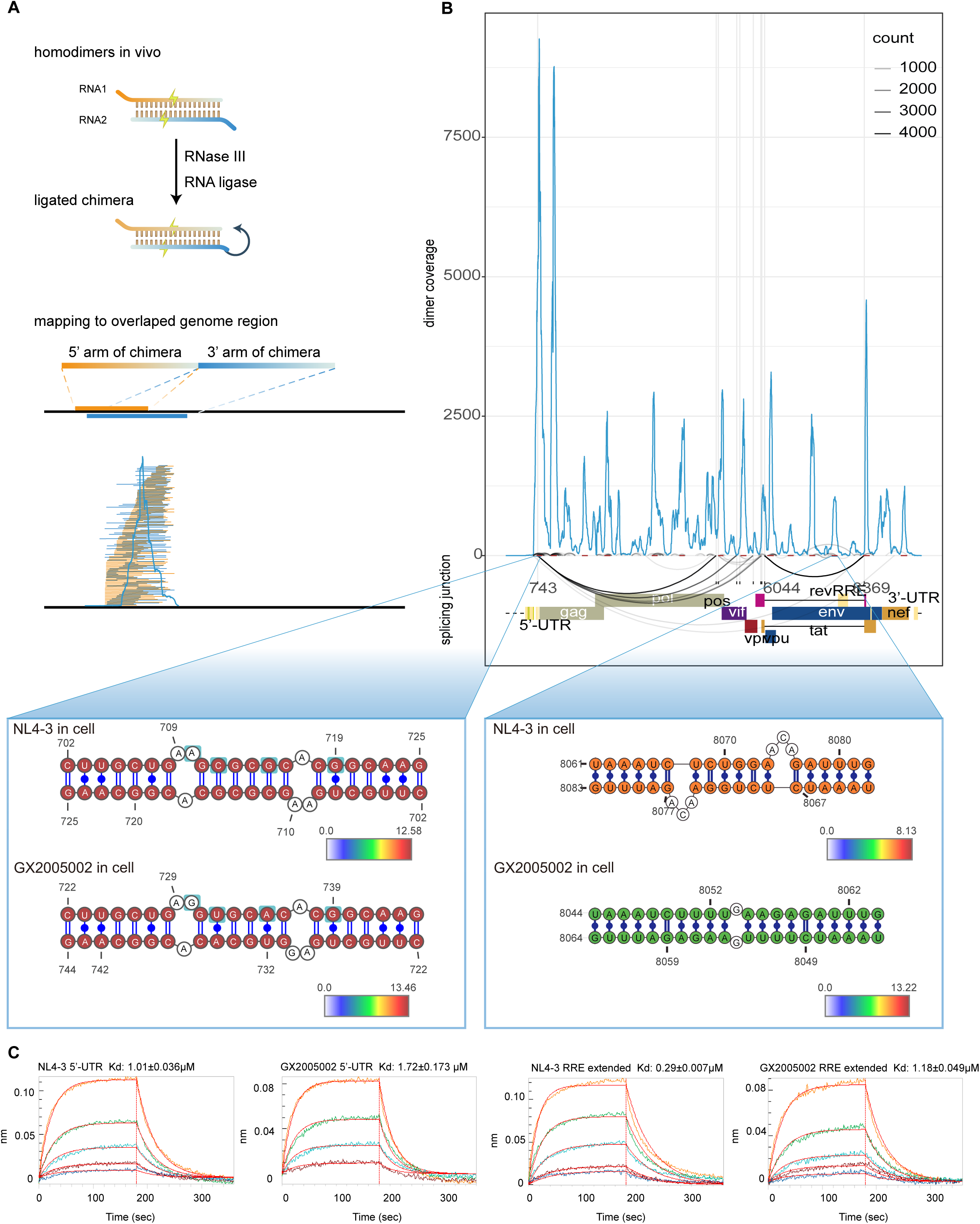
Identification and validation of homodimers in HIV-1 genome. A) Visual depiction of the RNA homodimer formation process: Inter-ligated fragments (arms) originating from homodimeric RNA molecules generate chimeras, where each arm aligns to overlapping coordinates on the HIV-1 genome. This process enables the plotting of coverage and specific details of dimers by utilizing the positions and counts of these overlapping chimeras. B) distribution of homodimers throughout the HIV-1 genome. The blue line plot showcases the homodimer coverage derived from ligated samples of NL4-3 infected cells along the HIV-1 genome. Arc plots exhibit discontinuous reads in non-ligated samples of NL4-3 infected cells, with dark red segments indicating peaks of homodimers. The lower panels depict the base pairing of homodimers in the SL1 region of the 5’-UTR and downstream of the RRE region. Color mapping indicates the log2 transformed dimer score. C) Assessment of dimer self-binding using Bio-Layer Interferometry (BLI). The data presented here are from three separate experiments, offering insights into how dimers interact and bind to themselves. The dissociation constant (Kd), indicated by the mean ± standard deviation, was determined from these experiments.

The strongest homodimerization signal was found in the 5’-UTR region, which aligns with previous studies. The 5’-UTR region is well-known for its role in triggering HIV dimerization. Previous literature has highlighted the importance of SL1 (DIS) and SL3 (Ψ) in 5’-UTR^8,31,32^. Our data shows that the SL1, SL2, and SL3 regions all have supportive dimeric chimeras (Supplementary Figure S7). We derived a Dimerization Score by calculating the reads that support base pairing of homodimeric chimeras (Supplementary Figure S8), analogous to the COMRADES Score described previously ^17,19,22^. The base pairing of homodimers in the SL1 region of the HIV genome is consistent with previous data from NMR and cyro-EM studies ^33,34^. Additionally, we observed that variations in nucleotides within the SL1 region do not alter the base pairing pattern of dimers between the NL4-3 and GX2005002 strains (Figure 3B) which is egret with previous structure robustness revealed by permutation and structure prediction results Strikingly, in addition to the known homodimer within the SL1 and SL3 regions, we observed homodimerization distributed along the entire length of the HIV genome (Supplementary Figure 9). Homodimers were present in both infected cells and virions of both strains, with approximately 20 peaks of dimerization that are conserved between the NL4-3 and GX2005002 strains along the genome (Suplementary table2).

We investigated whether dimers could be predicted by the strength of base pairing of intermolecular loop-loop interactions by performing a systematic molecular hybridization analysis on the genomes of two HIV-1 strains, NL4-3 and GX2005002. Our findings showed no correlation between the predicted folding energy and the abundance of measured dimeric chimeras (Supplementary Figure S9A), suggesting that the formation of HIV homodimers is not solely a consequence of local base pairing propensities. Instead, it implies that additional factors, such as the binding of specific proteins, may significantly influence the dimerization process, potentially playing a more decisive role in stabilizing or facilitating the formation of these homodimers.

The identification of multiple dimers in HIV RNA beyond the 5’-UTR region raises intriguing questions regarding their potential roles serving within the HIV genome. To this end, we analyzed the sequence within dimer peaks, identifying a prevalent AG-rich motif rich, present in nearly every peak. This motif highly? resembles the RNA binding motif of Serine/Arginine-Rich Splicing Factor (SRSF) (Supplementary Figure S9B,S9C and S10), which are essential RNA-binding proteins involved in pre-mRNA splicing and alternative splicing regulation^35^, serving as the main type of HIV-1 splicing factors^36^. This result suggests a potential role of dimerization in RNA splicing processes within the HIV genome.

5’-3’ discontinuous reads were identified in non-ligation control data, enabling the inference of splicing junction sites. These inferred sites showed strong concordance with canonical splicing sites and recent nanopore sequencing data^37^ (Figure 3B). Notably, almost every junction site, including splicing donor and acceptor sites, exhibited dimer peaks around them. Additionally, the majority of dimeric chimeras covered these junction sites and enriched downstream of acceptor sites (Supplementary Figure S10), suggesting that dimerization process/events takes place on unspliced genomic RNA. Interestingly, dimers surrounding splicing regulatory elements (as summarized in^36^) form stable base pairing supported by numerous chimeras (Supplementary Figure S10). This observation underscores the close proximity and potential functional relationship between dimerization events and RNA splicing processes at these critical sites within the HIV genome, suggesting a potential role of dimerization in splicing processes or assisting in the transport of unspliced RNA out of the nucleus.

According to a previous report that utilized CLIP-seq to analyze the RNA binding specificity of the Gag protein in both cells and virions/From a previous CLIP-seq analyzing of Gag protein’s RNA binding specificity, it was observed that Gag exhibits a preference for binding to multiple elements, such as psi and RRE, in HIV infected cells^38^. Interestingly, significant dimer peaks were also identified in these regions (Supplementary Figure S9A). This suggests a potential connection between Gag binding and dimerization of the HIV genome.By plotting contact matrices derived from non-overlapping and dimeric chimeras around the RRE region, we were able to identify two distinct dimer peaks flanking the RRE region (Supplementary Figure S11A).

To validate these novel homodimers, we synthesized extended RRE RNA fragments with two dimer sites using in vitro transcription and confirmed their ability to form a dimeric conformation through annealing and non-denaturing gel electrophoresis (Supplementary Figure S12A), as well as Agilent Tapestation 4200 capillary electrophoresis (Supplementary Figure S12B) and Bio-Layer Interferometry (BLI) technology (Figure 3C).

### 4. Interplay between dimeric peaks and 3D genome organization

The local interaction surrounding the RRE forms an extended "arch" structure that encompasses this element, as illustrated in Supplementary Figure S13. This unique architecture may contribute to stabilizing the core RRE conformation, providing a structural basis for the rev-RRE interaction and HIV RNA transport. Additionally, we observed substantial local interaction signals around dimer sites in the 5’-UTR, as depicted in Supplementary Figure S7. To provide a comprehensive overview of the local interaction landscape around dimer sites, we generated meta matrices by computing local interactions (from non-overlapping chimeras) at dimer sites and the flanking regions located 1/2 upstream and 1/2 downstream (Supplementary Figure S12). This analysis unveiled significant local interactions around dimer sites, suggesting the involvement of stable local structures in the formation of dimers between two copies of the HIV genomes.

We observed the meta matrices around dimer sites are similar to genomic domains, which have been previously described in SARS-CoV-2^22^. This observation prompted us to investigate the potential existence of genomic domains in the HIV RNA genome. Using similar methods^39^, we calculated insulation scores for two strains of the virus in both cells and virions, as depicted in Figure 4A.

**Figure 4.**
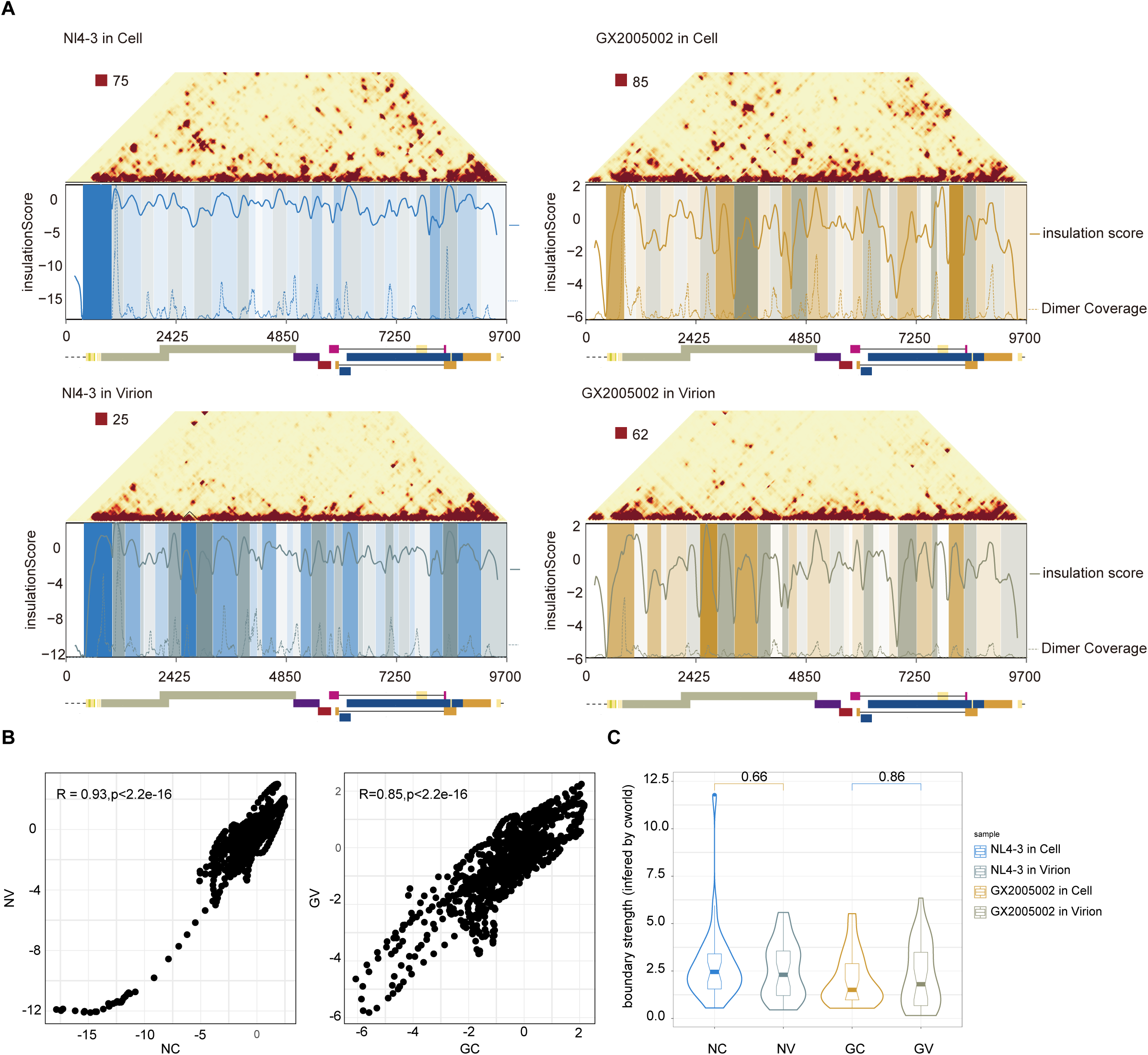
genome domains along HIV-1 genome. A) Each panel shows triangle matrix and genome domains of two strains of the virus in both cell and virion states, as calculated using C-world. The x-axis represents the genomic position, while the y-axis represents the insulation score (solid lines) and dimer coverage (dashed lines). B) Correlation of insulation score in infected cells and virion for NL4-3 and GX2005002 strain. C) Violin and boxplot comparing the boundary strength of the genomic domains of two strains between infected cells and virions. The boxplot displays the median, quartiles, and range of the boundary strength values for each strain in each state.

In this way, we identified 31, 33 HIV genomic domains in infected cells and 36, 32 domains in virions for NL4-3 and GX2005002 strains, respectively. The high correlation of insulation scores between cells and virions and consistent domain boundaries are observed (Figure 4B), and more importantly, the boundary strengths did not show any significant differences between cells and virions (Figure 4C), suggesting conserved, stable and persistent genomic domains during the analyzed stages.

Interestingly, we found that dimer sites are often located within genomic domains, and overlaying dimer coverage onto domains and insulation score curves revealed a striking concordance between dimer sites and genomic domains (Supplementary Figure S14). These findings suggest an interplay between the dimerization signals and global 3D organization of the HIV genome, providing insights into the complex mechanisms of HIV-1 replication.

### 5. Dynamic changes in global and long-range interactions throughout HIV-1 life cycle

We noticed extensive long-range interactions between the HIV-1 genome of two strains (NL4-3 and GX2005002) (Figure 1B and Supplementary Figure S4).

To quantitatively analyze these interactions, we utilized DESeq2, following a similar approach as previous studies. Our results showed that both strains exhibit a substantial number of long-range interactions in infected cells, while these interactions are significantly reduced in virions (Figure 5A). Furthermore, histograms of enriched interaction pairs revealed a significant decrease in long-range interactions exceeding 2500nt in virions compared to cells (Supplementary Figure S15A). The Contact probability decay curves (PS curve) also demonstrated a faster decay of genomic interactions in virions compared to cells, for both strains of the HIV-1 virus (Supplementary Figure S15B).

**Figure 5.**
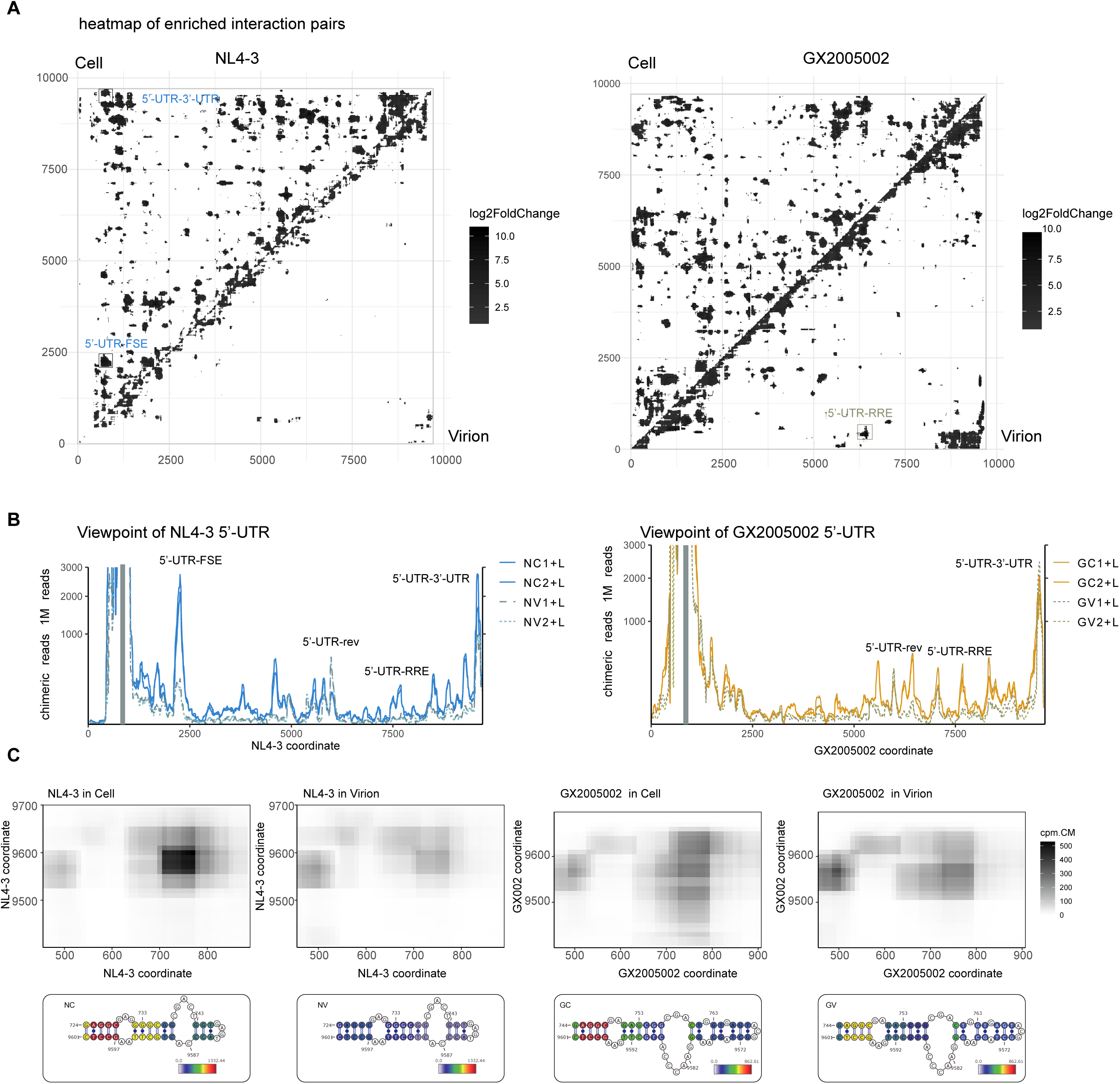
long range interaction in HIV-1 genome. A) heatmaps of enriched interactions obtained from NL4-3 and GX2005002 infected cells and virions. The upper diagonal shows interactions from infected cells, while the lower diagonal region displays interactions from virions B) Viewpoint lines depicts the binding positions of the 5’-UTR along the HIV-1 genome. The gray rectangles indicate the viewpoint regions. The colors of the lines represent specific samples, with samples from virions shown as dashed lines. C) Contact matrices and base pairing details between 5’-UTR and 3’-UTR. The top panels display heatmaps indicating contact probability, with the color bar indicating chimeric reads per 1M reads in each specific sample. The bottom panels show base pairing colored by base pairing scores.

Specifically, we employed viewpoint analysis to examine interactions involving the 5’-UTR. The results, depicted in Figure 5B, reveal multiple peaks of 5’-UTR interactions across the HIV-1 genome. Interestingly, for the majority of these peaks, the signals are lower in virions compared to cells. However, cyclization interactions (5’-UTR-3’-UTR) at the ends are still maintained in virions (Figure 5C). Intriguingly, we also identified multiple interaction peaks located near crucial elements, such as the frameshift element (Supplementary Figure S16A), rev start codon (Supplementary Figure S16B), RRE, and nuclear exporting signal (NES) (Supplementary Figure S16C), indicating complex roles of 5’-UTR in HIV life cycle. Among these interactions, some are strain specific, such as the interaction between 5’-UTR and FSE is specific to the NL4-3 strain, while interaction between 5’-UTR and 8.5K region (near NES) are highly enriched in GX2005002 strain (Supplementary Figure S16C). As mentioned earlier, the majority of interactions decrease in the virion, with the notable exception of interactions in the 5’-UTR and 6K region (near D4 splicing donor site), which significantly increase in the virion. (Supplementary Figure S16B). The enhanced interactions in virions are consistent across both strains.

These results indicate that the HIV genome undergoes systematic remodeling during virus packaging, with a general loss of long-range interactions but maintenance of specific interactions, including genome cyclization. The exploration of long-range interactions and their dynamics has provided us with an unprecedented understanding of the structural organization of the HIV genome. The significance of these interactions and their impact on the HIV life cycle and pathogenesis warrants further investigation in future research.

## discussion

### 1. HicapR : a reliable and efficient method for exploring structures of viral RNA genomes

In this study, we have introduced the HiCapR protocol, which combines the SPLASH method with the capture of HIV RNA-derived libraries. This integration of RNA extraction, fragmentation, proximity ligation, library construction, and subsequent capture of target RNA involved in interactions enables the investigation of various viral genomic structures. This protocol offers a streamlined and efficient approach for studying the structure and dynamics of complete HIV RNA genomes.

HiCapR uses a similar hybridization-based capture strategy as COMRADES^17,19^ except that in HiCapR, this capture step occurs after library construction, whereas in COMRADES, it takes place prior to proximity ligation. The post-library-capture principle employed in HiCapR is a well-established approach utilized in other methods such as capture Hi-C or capture-C^40-43^, and it offers additional flexibility relative to previous protocols. Our results demonstrate remarkable sensitivity and reliability of the HiCapR technique in capturing low-abundance HIV RNA structure, providing an unprecedented resolution and comprehensive approach to elucidate the intricate RNA structures within the HIV-1 genome. This novel proximity ligation method transcends the limitations of traditional techniques, offering a robust framework for dissecting the complex architecture of viral RNA.

### 2. New insight of HIV RNA local and long-range interactions

The 5’-UTR is crucial for various stages in the HIV life cycle. Our analysis of its structures across different strains and conditions revealed consistent canonical stem-loops and interactions, despite sequence variations and alternative conformations, highlighting the structural robustness? stability and functional significance of the HIV 5’-UTR.

In addition to its local folding, the 5’-UTR engages in extensive and dynamic long-range interactions, notably with the 3’-UTR, suggesting genome cyclization akin to other viruses (Figure 4). This phenomenon, also observed in Zika, influenza, and SARS-CoV-2^15,17,19,22^, hints at broader biological implications warranting further exploration. Additionally, the 5’-UTR interacts with key import elements in the HIV genome, such as NES, RRE, and the rev start codon. Given the Rev protein’s roles in RNA transport and nuclear export^44^, the 5’-UTR may also contribute to these processes.

Previous studies have highlighted folding principles of Zika, SARS-COV-2 coronavirus, and influenza virus, unraveling the intricate mechanisms underlying viral replication, pathogenesis, and host interactions ^15,17,20,22,45^. One of the key findings from this study is that the HIV-1 genome is organized in a complex three-dimensional structure that facilitates long-range interactions between distant regions of the genome. This study shed light on the dynamics of long-range interactions between the HIV-1 genome of two strains, NL4-3 and GX2005002. These interactions are largely reduced in virions compared to cells, suggesting a critical role in virus assembly and release.

Previous studies has elucidated the folding principles of various viruses like Zika, SARS-CoV-2, and influenza, shedding light on viral replication, pathogenesis, and host interactions ^15,17,20,22,45^. Our study revealed that the HIV-1 genome’s intricate three-dimensional organization enables long-range interactions.

The contrasting loss of long-range interactions in the HIV-1 genome compared to SARS-CoV-2 ^22^ suggests distinct folding processes between these viruses. This disparity underscores the unique characteristics of each virus and their genomic structures. Speculations indicate that the HIV-1 genome may adopt a rod-like structure within viral particles, unlike the spherical compression seen in SARS-CoV-2 ^46^. Further investigation is needed to unravel the mechanisms and biological significance of this compression in the HIV-1 genome.

### 3. Emerging Insights into HIV-1 Dimerization through newly identified dimer sites

Our detailed analysis has consistently identified approximately 20 candidate dimer peaks both within and beyond the 5’-untranslated region (5’-UTR) across the NL4-3 and GX2005002 strains of HIV. Intriguingly, these peaks show a notable enrichment in the vicinity of splicing sites, frequently featuring a sequence motif that bears a strong resemblance to the RNA binding signature of the Serine/Arginine-Rich Splicing Factors (SRSF). This enrichment pattern and the presence of the SRSF-like motif at these sites suggest a potential regulatory role of these dimers in the complex landscape of HIV RNA splicing. The pervasive presence of homodimers flanking almost every splicing site introduces the intriguing hypothesis that these dimers could be actively participating in the regulation of alternative splicing or facilitating the export of unspliced genomic RNA from the nucleus to the cytoplasm. This hypothesis warrants rigorous investigation in subsequent studies to elucidate the mechanistic underpinnings of these observations. Collectively, these findings hint at a significant interplay between RNA dimerization and splicing processes within the HIV genome, which could have profound implications for our understanding of viral gene expression and replication strategies.

Moreover, significant dimer signals were observed in the RRE flanking sequences, adding complexity to its functional role, potentially impacting nucleocytoplasmic transport, Gag binding, and virus packaging processes. Given previous studies highlighting Gag binding to RRE and the crucial role of RRE in anchoring Gag synthesis^3,38,47^, it is further hypothesized that HIV-1 homodimers may play a role in HIV splicing or assist in transporting unspliced full-length RNA out of the nucleus for genome packaging.

Interestingly, the observation that candidate dimer sites are located within genomic domains suggests that dimerization may be influenced by the local RNA folding environment (Supplementary Figure S12). It is possible that the dense local interaction around these sites may facilitate dimerization by bringing the two regions of the genome into close proximity. Alternatively, dimerization may play a role in shaping the local chromatin environment by promoting even initiating the formation of genomic domains. Further research is needed to fully understand the relationship between RNA genomic domains and dimer sites in the HIV-1 genome. However, the identification of these candidate dimer sites within genomic domains provides a starting point for investigating the role of local genome RNA interactions in HIV-1 replication and dimerization. These findings may have implications for the development of novel antiviral therapies that target the dimerization process and may provide new avenues for future research in this field.

In summary, our study provides a comprehensive analysis of the HIV-1 genome using reliable HicapR, revealing potential dimer sites and long-range interactions, particularly within the 5’-UTR as well as persistent structure domains? These findings significantly advance our understanding of the structure, dynamics and robustness of HIV RNA and their involvement in splicing, assembly and packaging processes, potentially leading to the development of novel antiviral therapies.

## methods

### Cell culture and virus infection

The MT4 cell lines (RRID:CVCL_2632) were cultured under specific conditions: RPMI 1640 medium (Gibco, USA) supplemented with 10% fetal calf serum (Gibco, USA), 100 U/ml penicillin, and 100 μg/ml streptomycin. The cells were maintained at a temperature of 37°C with 5% CO2 and saturated humidity. To initiate infection, cells were exposed to HIV-1 NL4-3 (HIV-1 strain of subtype B) or HIV-1 GX2005002 (HIV-1 primary strain of CRF01_AE which is one of the main strain in China, accession: GU564222) at a multiplicity of infection (MOI) of 0.15.. Concurrently, parallel control groups consisting of uninfected cells were also established. Both the cells and the cell supernatant were collected after 48hours post infection for subsequent experiments.

### HicapR method

First, extracted crosslinked RNA was treated as in simplified SPLASH protocol ^22^.

Briefly, 500 ng of each sample was fragmented using RNase III (Ambion) in a 20 μl mixture for 10 minutes at 37°C. The fragmented RNA was then purified using 40 μl of MagicPure RNA Beads (TransGen). Each RNA sample was subsequently divided into two halves, with one half used for proximity ligation and crosslink reversal (C, V samples). The proximity ligation process was then carried out using the following conditions: 200 ng of fragmented RNA, 1 unit/μl RNA ligase 1 (New England Biolabs), 1× RNA ligase buffer, 1mM ATP, 1 unit/μl Superase-in (Invitrogen), with a final volume of 200 μl. The reactions were incubated for 16 hours at 16°C and were stopped by cleaning with the miRNeasy kit (Qiagen). To reverse the crosslinking, the RNA was irradiated on ice with 254 nm UltraViolet C radiation for 5 minutes using a CL-1000 crosslinker (UVP). For the non-ligated controls, crosslink reversal was performed immediately after crosslinking, without proximity ligation (these controls were labeled as "-L").

The proximity ligated RNA was then subjected to library construction using SMARTer Stranded Total RNA-Seq Kit v2 (Clonetech).

The cDNA libraries were enriched for HIV fragment using the TargetSeq One Hyb & Wash kit (igenetech) with the T548XV1 probe panel, designed based on HIV genome. The probe sequences are provided as supplementary data

### Data preprocessing and chimeric reads identification

The data preprocessing and identification of chimeric reads were conducted as previously described. The reference genome used for NL4-3 strain was a combination of NL4-3 genome sequence (https://www.ncbi.nlm.nih.gov/nuccore/AF003887) and human spliced mRNAs and noncoding RNAs described in ^30^, while for GX2005002 strain, the reference genome used was a combination of GX2005002 genome sequence (https://www.ncbi.nlm.nih.gov/nuccore/KP178420) and human spliced mRNAs and noncoding RNAs as above.

First, we used pear to merge the overlapped reads:

pear -e -j 32 -f sample_R2.fastq.gz -r sample_R1.fastq.gz -o Sample.PEAR note that for Clonetech 634413, the sense strand of RNAs is in R2 read.

Then, we used fastp to filter reads with low quality and cut adapters:

fastp -i sample.PEAR.fastq -o sample.PEAR.fastp.fastq -h fastp.html -w 4 -a AGATCGGAAGAGCGTCG

And then, chimeric reads are detected using hyb pipeline with default setting:

hyb detect in= sample .PEAR.fastp.fastq db=$DB qc=none

### Chimeras for RNA-RNA interaction and dimer identification

Homodimers were identified in accordance with the methods previously reported in the literature, with a more stringent filter^23^. Specifically, we utilized the hub pipeline and filtered chimeras in .hyb files. To calculate the degree of overlap between the two arms of chimeric reads, we used the formula L=1+min(e1,e2)-max(s1,s2), where e1 and e2 represent the ends of each arm of chimeric reads, while s1 and s2 represent their starts. To ensure that possible circular RNAs, which may be produced by single RNA end-to-end ligation, are filtered out, we applied the following condition: (e1 < e2) OR (s1 < s2). A schematic diagram of dimer chimeras is provided in Supplementary Figure S1.

We defined the dimer range as the maximum value between s1 and s2, and the minimum value between e1 and e2. Based on this range, we further calculated the coverage of dimer chimeras.

On the other hand, only non-overlapping chimeras (with L<0) were included for RNA-RNA interaction contact matrices to reduce the possibility that the chimeras called for RNA-RNA interactions come from homodimers.

### Dimer score and dimer base pairing

The dimer score (DS) was defined as the number of chimeric reads that supported a potential dimer base-pairing event. To visualize each particular candidate dimer, we used the hybrid-min command in the RNA fold package to in-silico hybridize two homodimers. The paired bases were then colored using the aforementioned dimer score.

### Local structure folding and MDS plot

Initial local structure folding was performed similarly to a previous report^17^. Base pairing scores were first calculated using comradesMakeConstraints function in COMRADES package(https://github.com/gkudla/comrades):

comradesMakeConstraints -i sample_rm_overlap.hyb -f Genome_fasta -b 1 -e genome_length

we folded the RNA structures 1000 times using the COMRADES manual. Subsequently, they utilized multidimensional scaling to calculate the distances between RNA structures.

### Calculation of base pairing probability for local structure

For a particular cluster in the MDS plot, we construct a base pairing probability matrix to reveal consensus stems in these structures, where the probability of base pairing for each base pair is determined by calculating the proportion of structures that support the base pair among all folded structures.

### Dimer validation

RNA preparation. 5’-UTR and RRE DNA fragments were amplified by RT-PCR using RNA from infected cells, with primers listed in Supplementary table3. Purified NL4-3 and GX2005002 5’-UTR and extended RRE PCR products (200 ng) were utilized as templates for RNA in vitro transcription with T7 RNA polymerase (Vazyme), following manufacturer’s protocol. The reaction was then incubated at 37 °C for 16 hours, followed by DNase I treatment for 30 minutes at 37 °C. The RNA was subsequently purified using MagicPure RNA Beads (TransGen) through gel purification.

### Native agarose gel electrophoresis

RNA (600 ng) was heated to 95°C and then slowly cooled to room temperature in either high salt buffer (50mM Tris-HCl pH 7.5, 140mM KCl, 10mM NaCl) or low salt buffer (high salt buffer diluted 1:10 with water). Samples were loaded with native loading dye (0.17% Bromophenol Blue and 40% (vol/vol) sucrose) on 2% agarose gel prepared with 1× tris-borate magnesium (TBM) buffer (89 mM Tris base, 89 mM boric acid and 2 mM MgCl2) and fractionated at 100 V for 85 min at room temperature. RNA in the PAGE gel was visualized using GelRed nucleic acid gel stain(Thomas scientific).

### Agilent 4200 TapeStation Capillary electrophoresis

We prepared the RNA samples as described above and loaded them onto the TapeStation RNA screentape without the heating and denaturing step. Subsequently, we initiated the device for electrophoresis analysis, which ran automatically without the need for manual intervention. After the electrophoresis analysis was completed, the system generated electropherograms and relevant data regarding the RNA samples, including their size distribution and concentration. This method allowed us to quickly and accurately evaluate the conformational features of the RNA.

### Biomolecular Binding Kinetics Assays

Biomolecular Binding Kinetics Assays were performed using the Octet R8 Platform (sartorius) following a standardized protocol. The assay involved the preparation of analytes at varying concentrations, real-time monitoring of association and dissociation phases, and subsequent data analysis to determine kinetic parameters kd. Quality control measures were implemented to ensure the reliability and reproducibility of the binding kinetics measurements.

## Data availability

The raw sequence data reported in this paper have been deposited in the Genome Sequence Archive ^48^ in National Genomics Data Center ^49^, China National Center for Bioinformation / Beijing Institute of Genomics, Chinese Academy of Sciences (GSA: CRA016024) that are publicly accessible at https://ngdc.cncb.ac.cn/gsa.

## Code availability

Custom code for the analysis performed in this study is publicly available via biocode at https://ngdc.cncb.ac.cn/biocode/tools/BT007456

## Supplementary Figure S1

This figure illustrates the workflow for identifying chimeric reads supporting RNA-RNA interactions as well as homodimers. The rounded rectangle in the upper right corner of the figure displays the start and end positions of the 5’- and 3’-arms of chimeric reads. These arms can be ligated in multiple ways to form chimeras, some of which are thought to be produced exclusively by homodimers.

First, sequencing reads are mapped to the HIV RNA genome, and chimeric reads are identified using hyb pipeline. Dimeric chimeras are then filtered to select only those that can be formed exclusively by dimerization. To reduce background noise, especially contaminated from homodimers, we selected non-overlapping chimeras for our analysis of RNA interactions.

This approach allowed us to identify both RNA-RNA interactions and candidate dimer sites and predict secondary structures with high coverage chimeras.

## Supplementary Figure S2

HIV Genome coverage of HiCapR data.

(A) The organization of the HIV NL4-3 genome follows NCBI annotation and is specified.

(B) and (C) The sequencing data from each group was mapped to the HIV genome and then genome coverage was calculated. The results showed that both NL4-3 (B) and GX2005002 (C) strains of the virus, whether in infected cells or virions, and whether RNAs are ligated or not, had uniformly complete genome coverage. This suggests that the HiCapR method is able to capture the entire HIV genome with low bias.

## Supplementary Figure S3

The heatmap displays the correlation between samples, which was measured using Pearson’s coefficients.

## Supplementary Figure S4

Related to Figure 1B, but with the inclusion of information from individual replicates and data from non-ligation controls, offering a more detailed view of the interactions and distributions presented in Figure 1B.

## Supplementary Figure S5

The established conformation of the HIV-1 5’-UTR, supported by chimeras in NL4-3 and GX2005002, both in cells and in virions. Colors of the nucleotides indicate the log2 transformed base pairing scores.

## Supplementary Figure S6

The stacked bar chart displays the number of 3’-5’, 5’-3’, and dimer chimeras in various ranges across the sample groups. The panel on the left pertains to NL4-3, while the one on the right pertains to GX2005002.

## Supplementary Figure S7

(A) Heatmaps simultaneously plot contact matrices calculated from non-overlapped chimeras, which is informative for interactions and dimer chimeras which is derived for homodimers. The coordination in heatplota is RNA-based, with the numbers in brackets indicating the corresponding coordinates in the DNA genome.

(B) Homodimer chimeras details around splicing sites show the 5’-arm represented by blue lines and the 3’-arm represented in yellow, providing a visual representation of the interactions in these regions.

## Supplementary Figure S8

Dimer score matrices around 5’-UTR of NL4-3 and GX2005002 strain in different stages.

## Supplementary Figure S9

(A) Homodimer coverage along HIV-1 genome in NL 4-3 and GX2005002 both in cells and in virion samples. Gag-CLIP data from Kutluay, S. B., et al. (2014) are also shown. Bottom panel depict delta G of each100-nt window in NL4-3 genome.

(B) Distribution of enriched motif in homodimer peaks of NL4-3, region similar to SRSF motif was underlined in the seqlog.

(C) Distribution of enriched motif in homodimer peaks of GX2005002, region similar to SRSF motif was underlined in the seqlog.

## Supplementary Figure S10

Top: Homodimer chimeras details around splicing sites show the 5’-arm represented by blue lines and the 3’-arm represented in yellow, providing a visual representation of the interactions in these regions. Known splicing regulatory elements are also indicated.

Bottom: basepairing of homodimers around splicing sites as indicated. Color code indicates log2 transformed base pairing score supporting each base-pair.

## Supplementary Figure S11

(A) Contact matrix and homodimeric chimeras surrounding the RRE region in the HIV genome. The left panel displays the contact matrix derived from non-overlapping chimeras and the contact matrix derived from dimeric chimeras. The right panel presents specific details of the dimeric chimeras around the RRE elements.

(B) Base pairing of homodimers in the extended RRE region in specific samples. Color mapping indicates the log2 transformed dimer score.

## Supplementary Figure S12

(A) Native agarose gel confirming dimeric conformation of RRE (extended to 620nt)

(B) Agilent tapestation 4200 capillary electrophoresis confirming dimeric conformation of 5’-UTR and RRE (extended to 620nt)

## Supplementary Figure S13

Predicted structure of extended RRE. Color code indicates log2 transformed base pairing score supporting each base-pair.

## Supplementary Figure S14

Average contact matrices around peak centers. Heatmaps showing the normalized average interaction frequencies for all peak centers as well as their flanking regions (±0.5 peak width). The heatmaps were binned at 10 nt resolution.

## Supplementary Figure S15

(A) histograms show distributions of different types of interactions across the HIV genome. The x-axis represents the span of the interactions, while the y-axis represents the frequency of each interaction type.

(B) Contact probability decay curves (Ps curves) showing progressive reconfiguration HIV genomes. Both two strains of the virus exhibit faster decay rates within virions than cells.

## Supplementary Figure S16

Contact matrices and base pairing details between 5’-UTR and 2.2K (A), 6K (B) and 7.6∼8.5K (C).

## Supplementary data

### Supplementary table1

Mapping statistics

### Supplementary table2

Homodimer peaks information

### Supplementary table3

Primers used for amplifying 5’-UTR and extended RRE

### Supplementary data

Designed probes for enriching HIV chimeras in fasta format

## Notes

### Competing Interest Statement

The authors have declared no competing interest.

### Summary of Updates

wo have majorly modified figure 1 and results part 1

