## Supplementary Figures for "Mapping HIV-1 RNA Structure, Homodimers, Long-Range Interactions and persistent domains by HiCapR"

Supplementary Figure S1

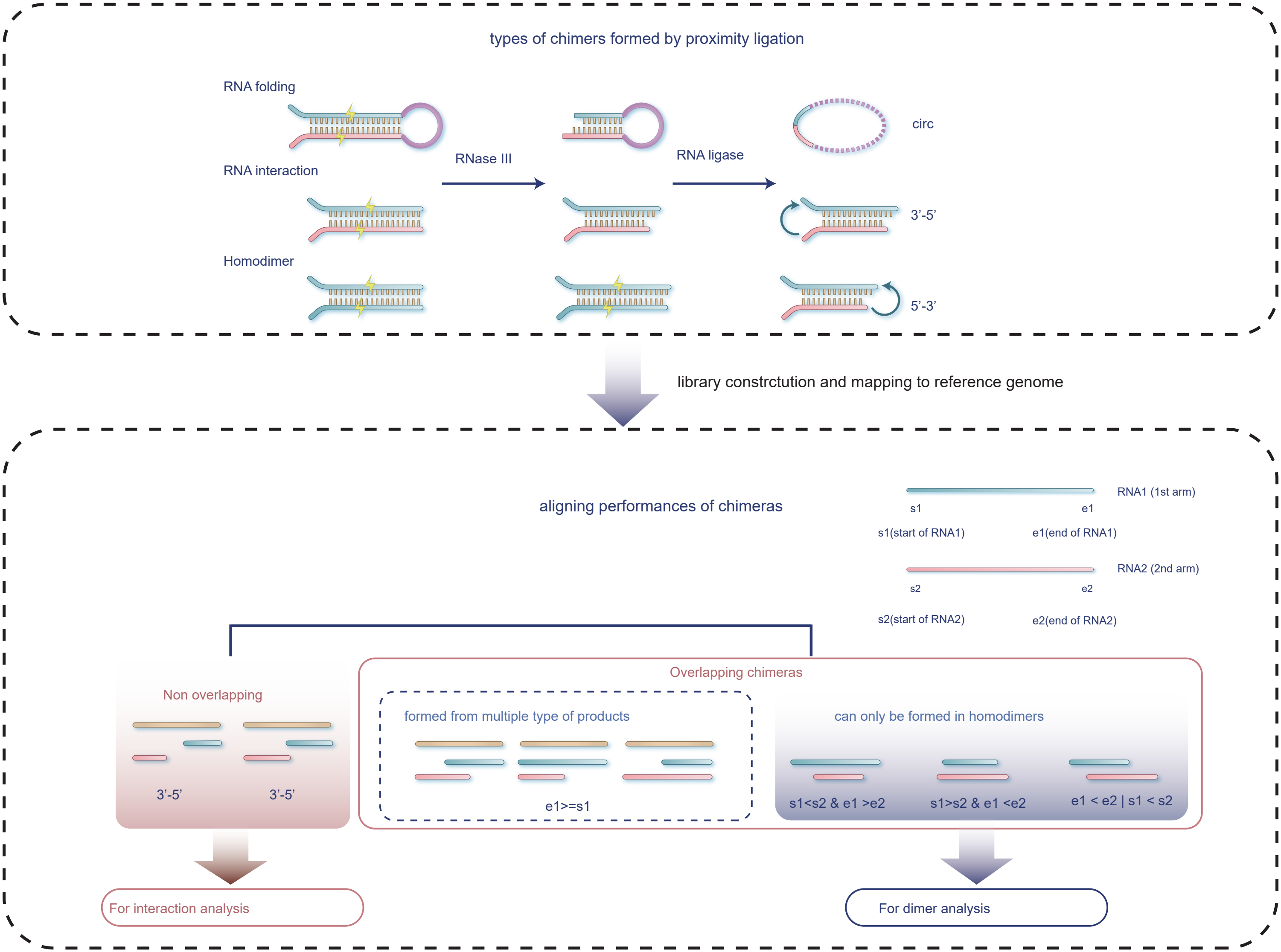

### Supplementary Figure S2

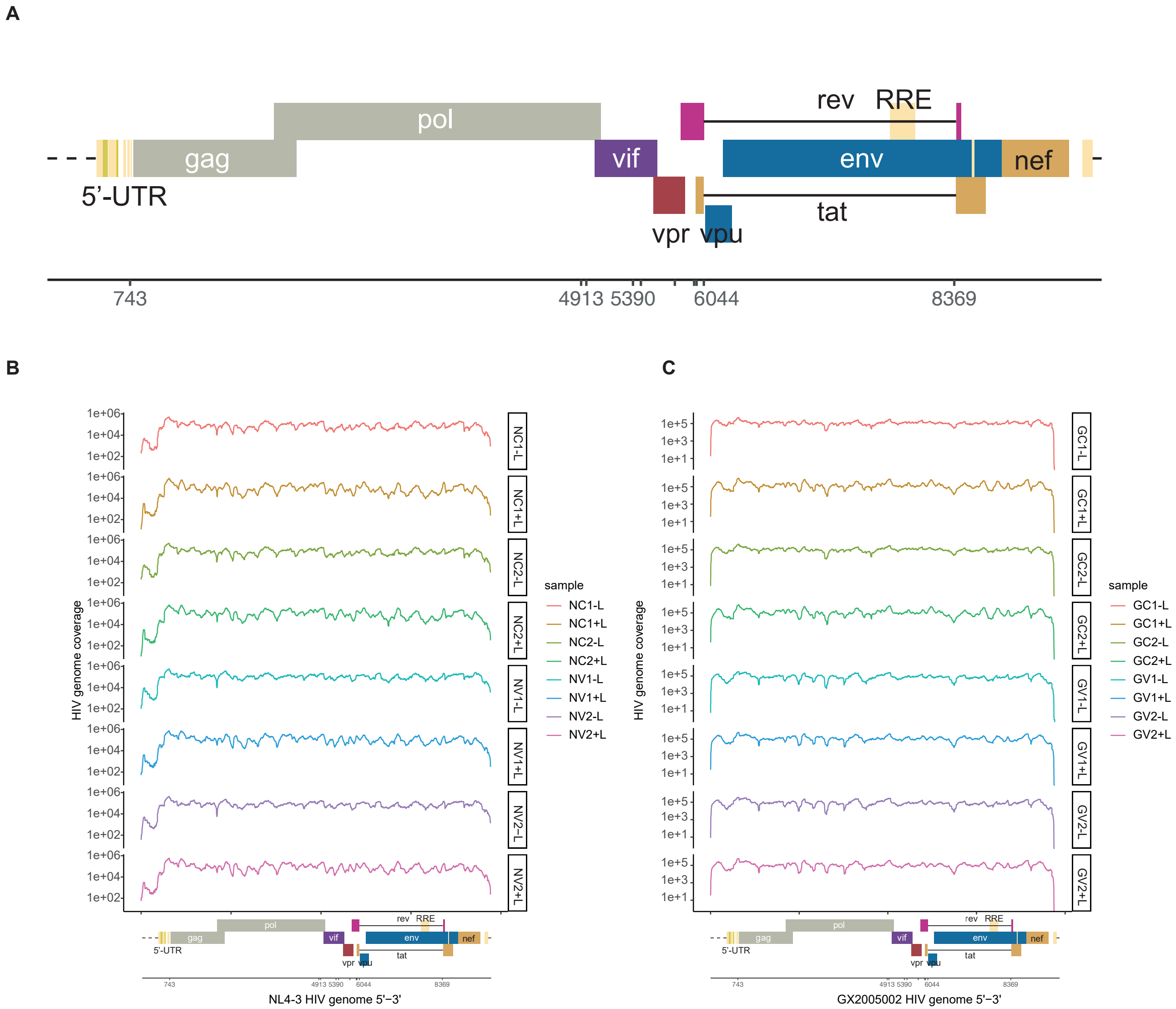

### Supplementary Figure S3

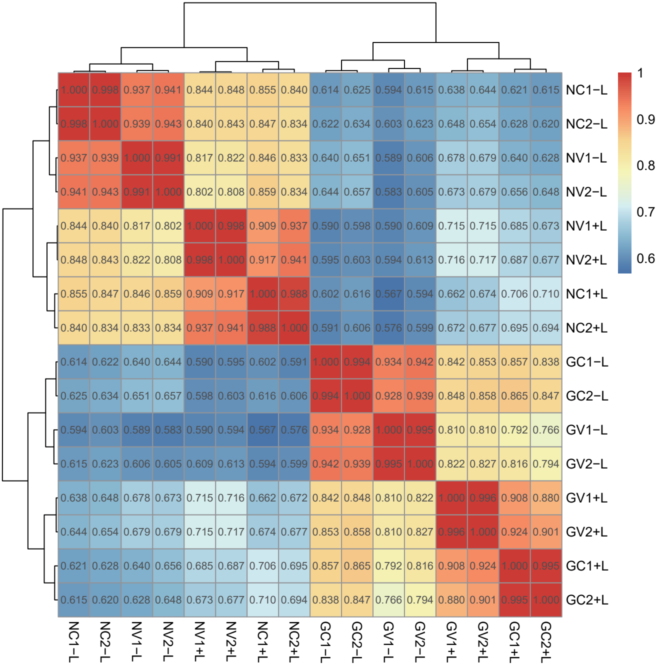

Supplementary Figure S4

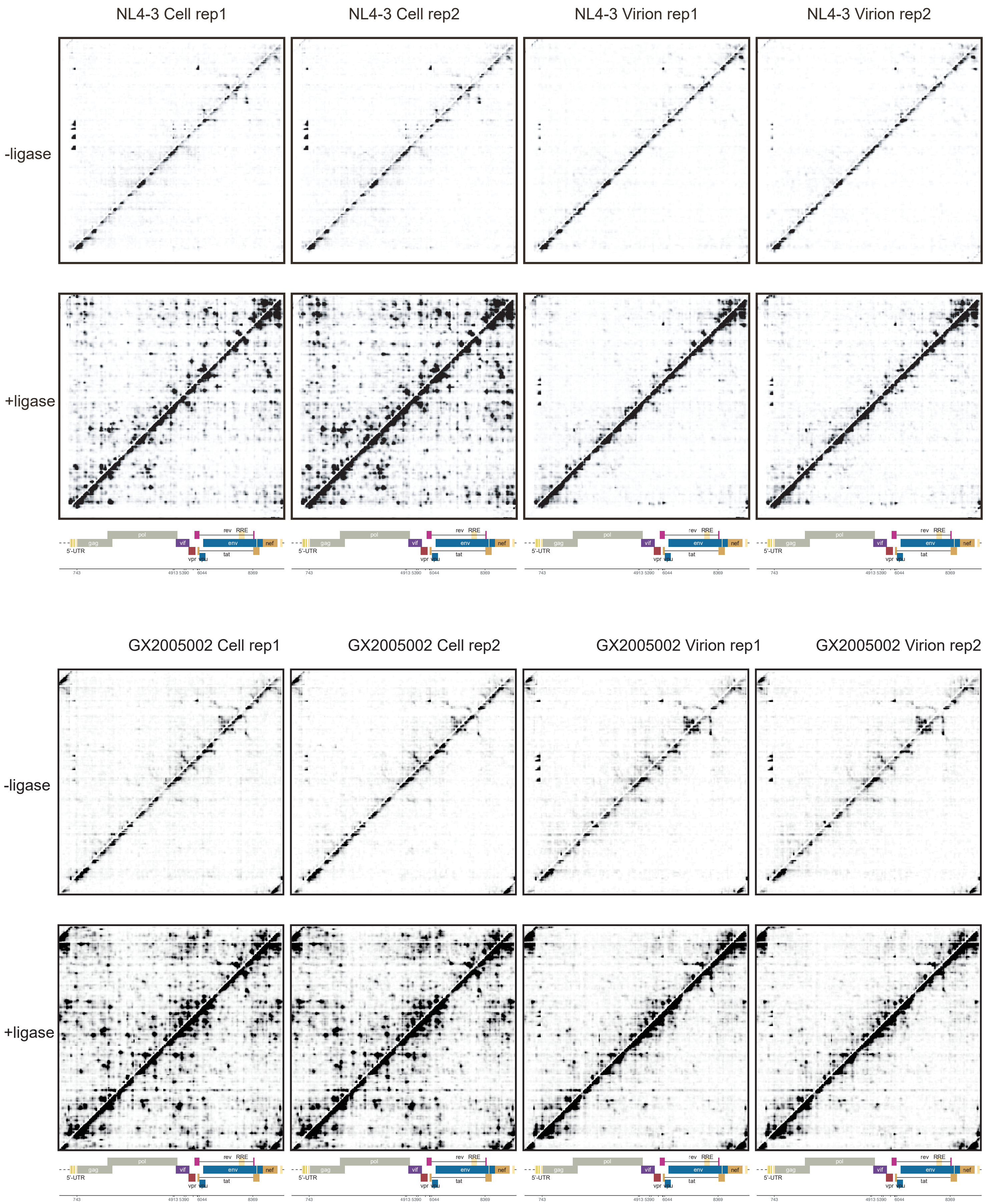

### Supplementary Figure S5

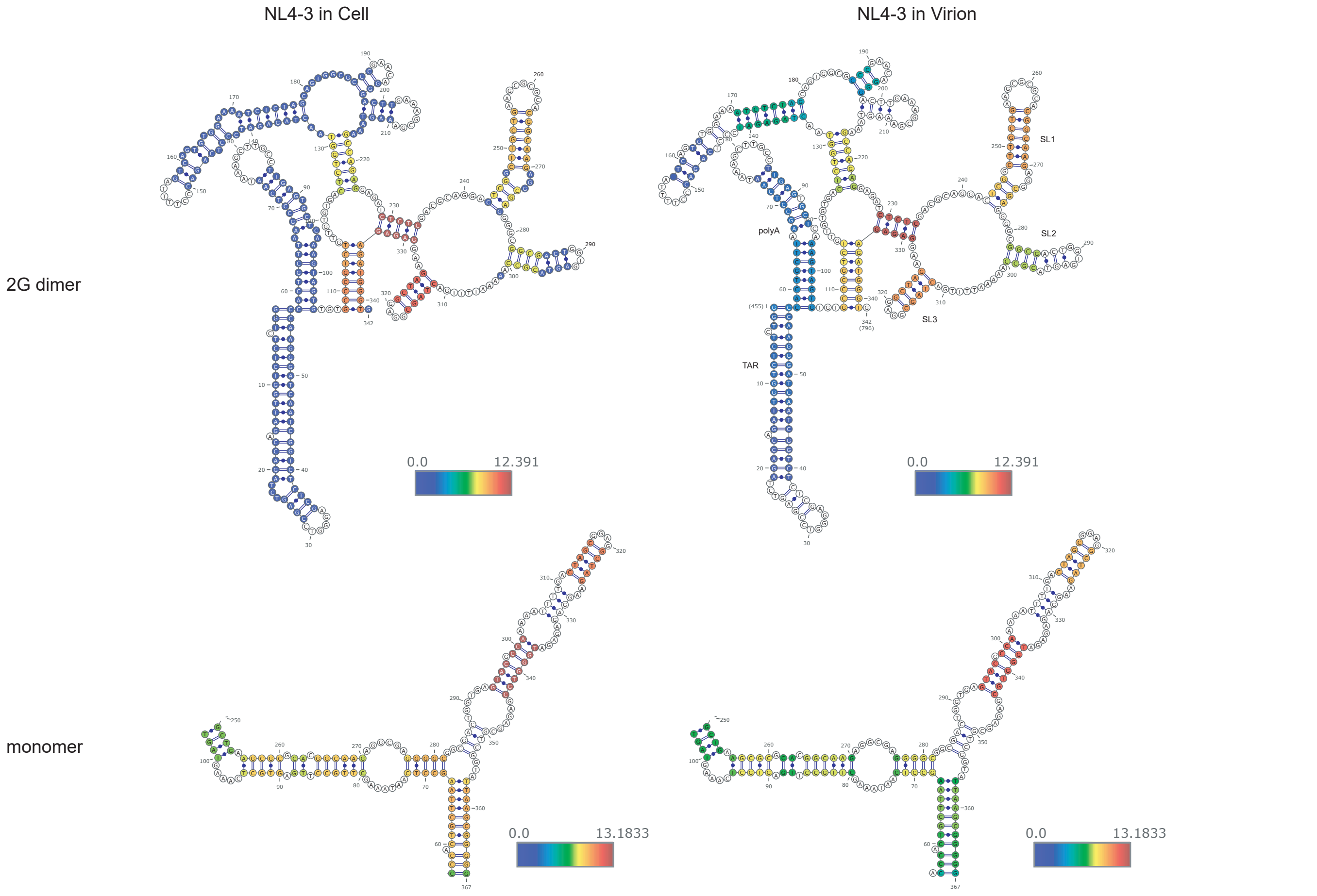

### Supplementary Figure S6

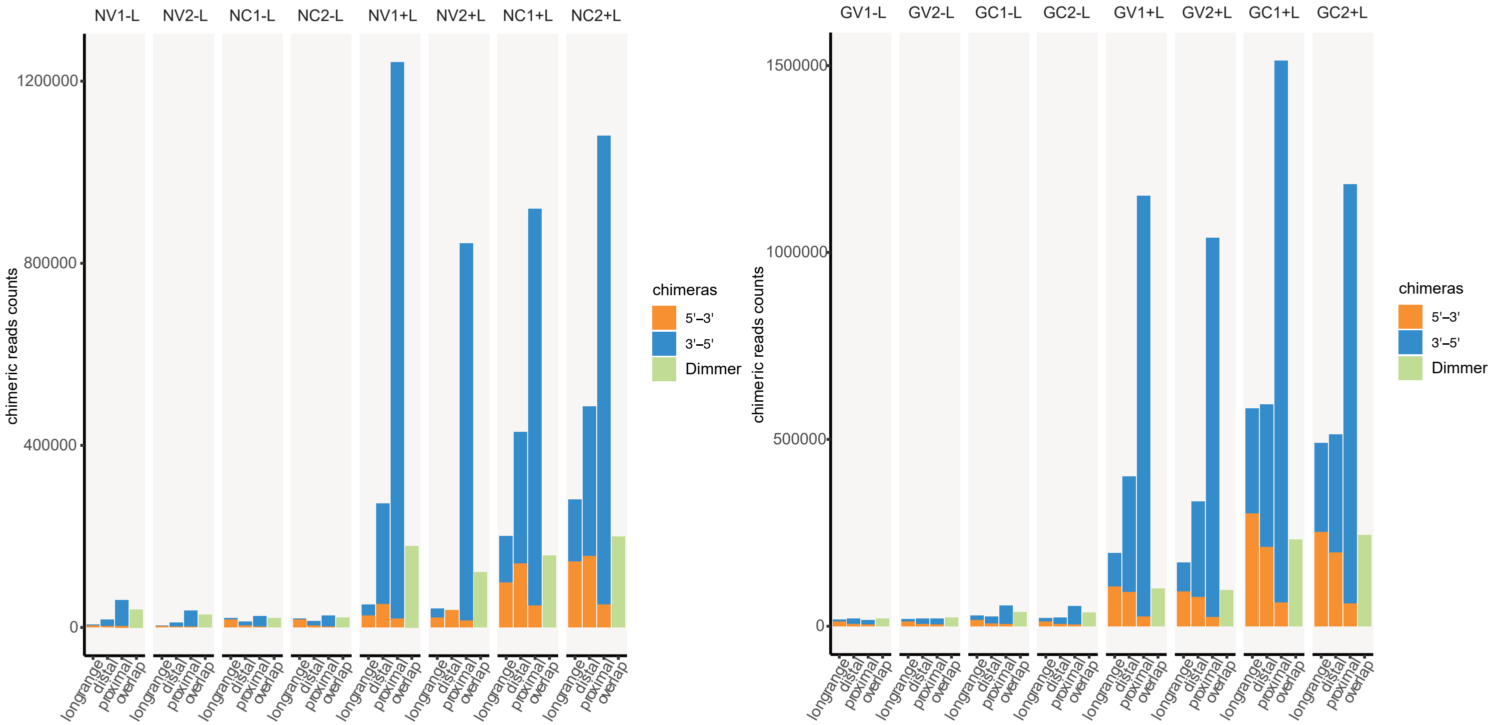

### Supplementary Figure S7

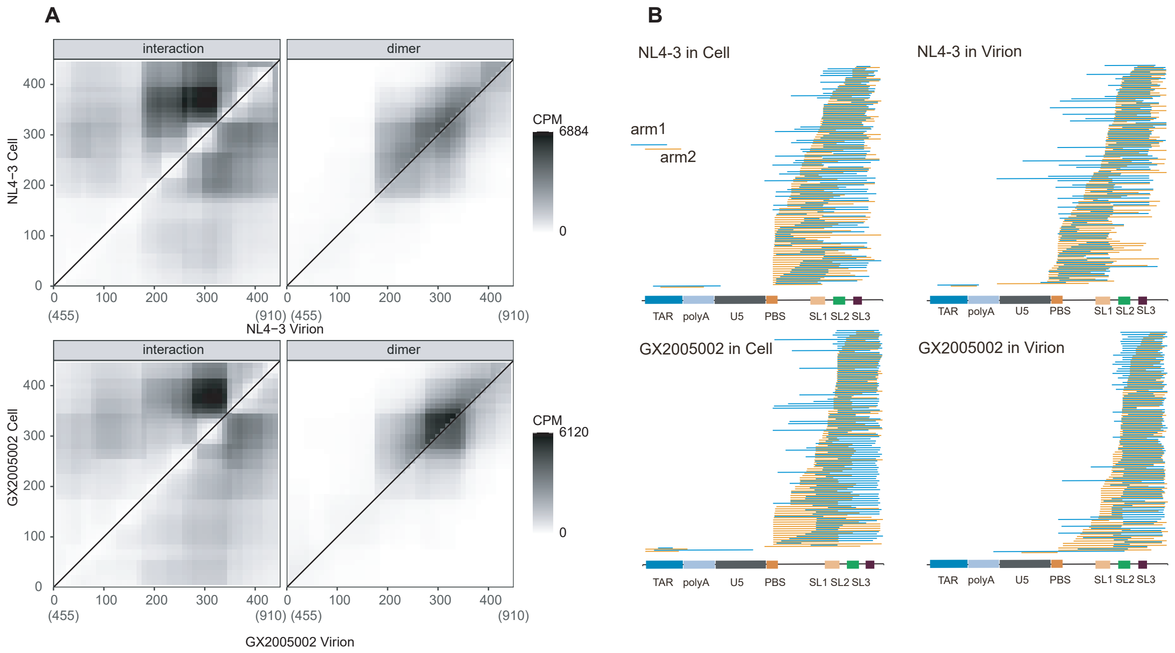

### Supplementary Figure S8

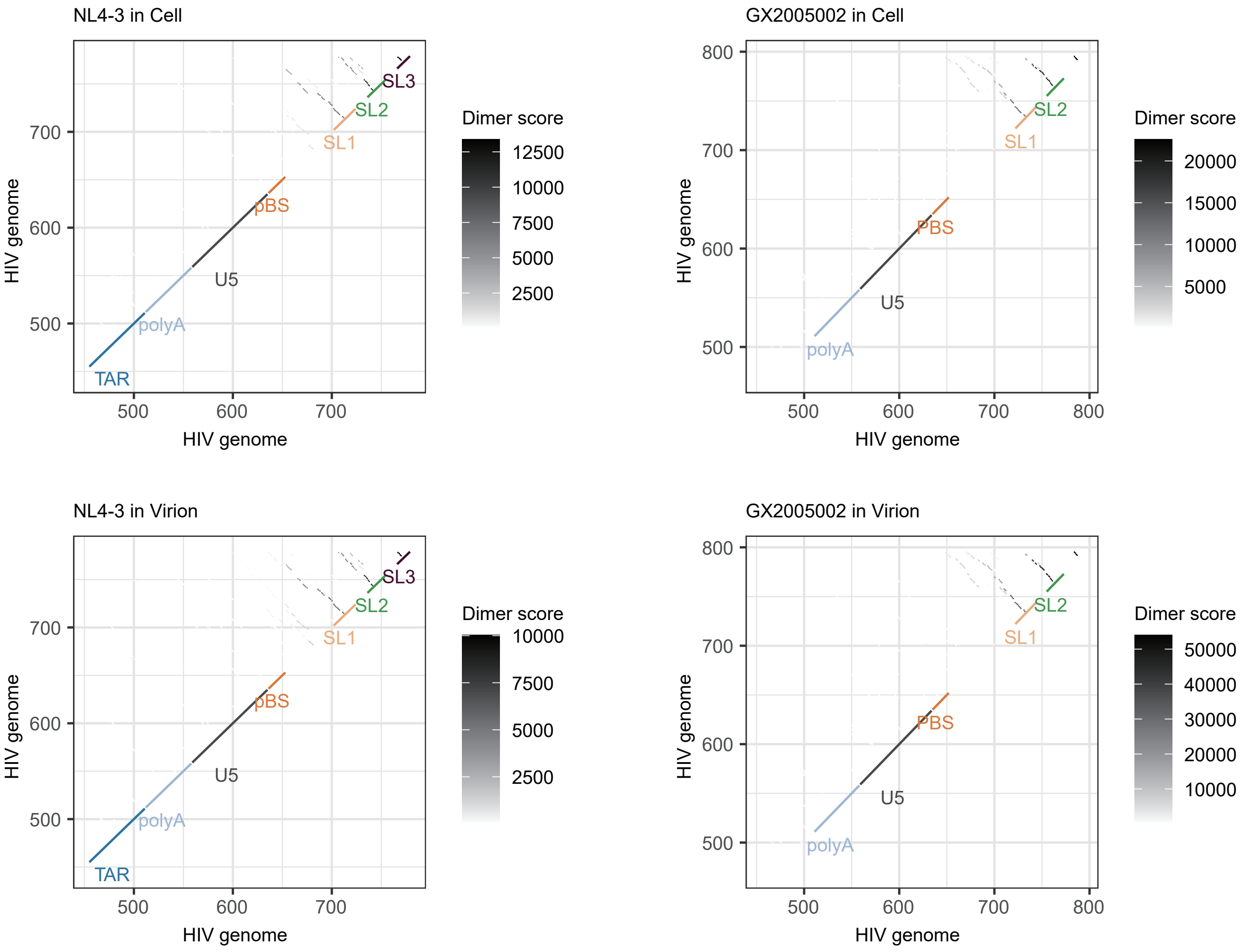

### Supplementary Figure S9

**A**

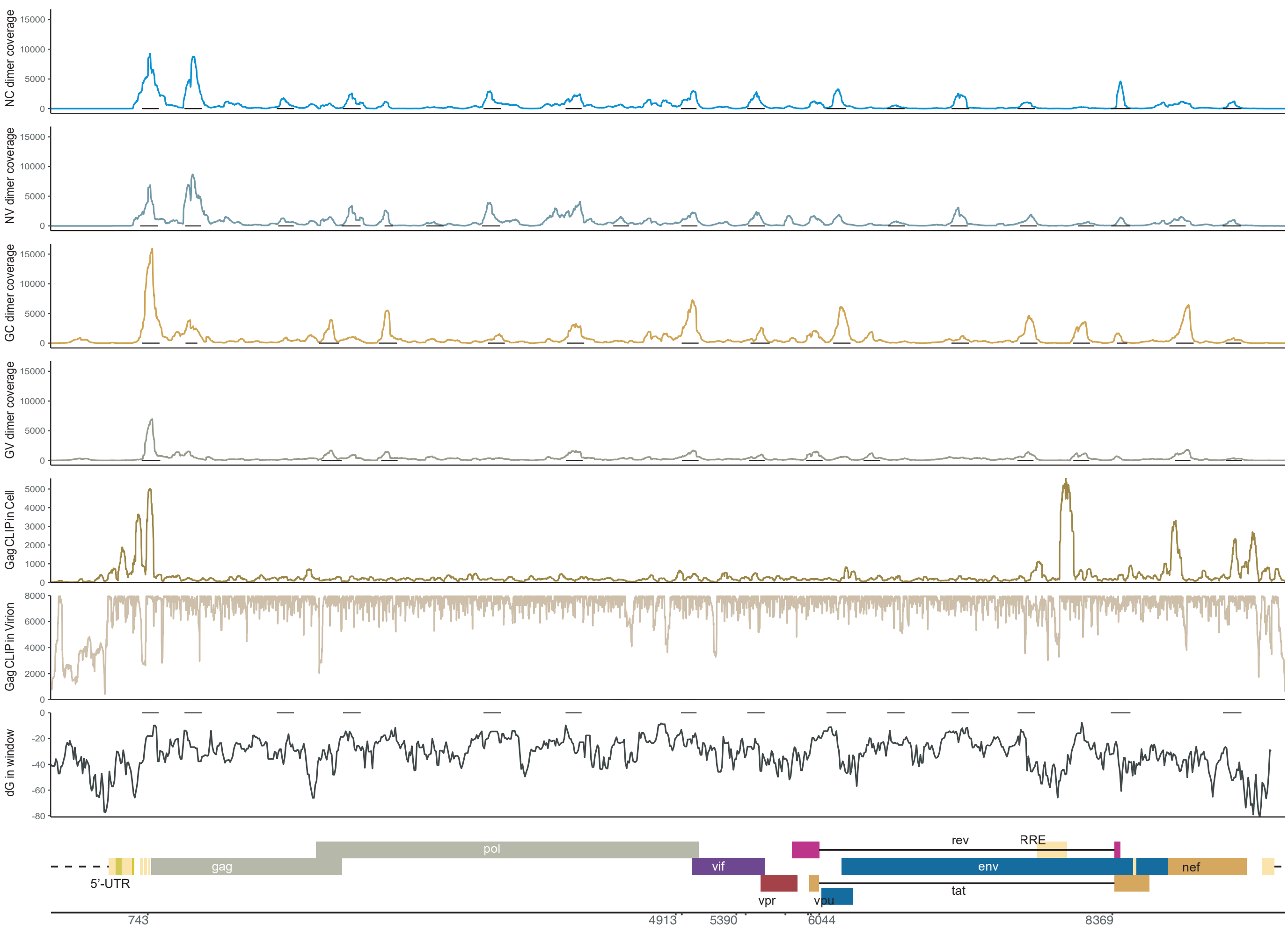


**B**

**SRSF motif**

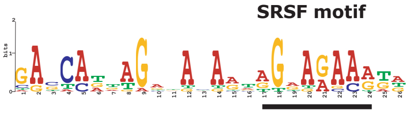

| | Name | Strand | Start | *p*-value | Sites | | |
|---|---|---|---|---|---|---|---|
| 3. | NL4-3:1778-1914 | + | 24 | 1.68e-10 | AAAGCATTGG | GACCAGGAGCGACACTAGAAGAAATG | ATGACAGCAT |
| 11. | NL4-3:7087-7223 | + | 71 | 4.34e-10 | GGACCAGGGA | GAGCATTTGTTACAATAGGAAAAATA | GGAAATATGA |
| 9. | NL4-3:6103-6256 | + | 88 | 1.98e-8 | TTAATTGATA | GACTAATAGAAAGAGCAGAAGACAGT | GGCAATGAGA |
| 13. | NL4-3:8339-8494 | + | 61 | 2.21e-8 | GGACCCGACA | GGCCCGAAGGAATAGAAGAAGAAGGT | GGAGAGAGAG |
| 6. | NL4-3:4050-4177 | + | 66 | 8.24e-8 | TCAAATAATA | GAGCAGTTAATAAAAAAGGAAAAAGT | CTACCTGGCA |
| 4. | NL4-3:2300-2439 | + | 32 | 1.20e-7 | TTAGATACAG | GAGCAGATGATACAGTATTAGAAGAA | ATGAATTTGC |
| 8. | NL4-3:5479-5617 | + | 98 | 2.06e-7 | AGCCCCAGAA | GACCAAGGGCCACAGAGGGAGCCATA | CAATGAATGG |
| 1. | NL4-3:717-850 | + | 104 | 2.45e-7 | TTAAGCGGGG | GAGAATTAGATAAATGGGAAAAAATT | CGGT |
| 7. | NL4-3:4958-5081 | + | 69 | 3.44e-7 | GAAAAGCAAA | GATCATCAGGGATTATGGAAAACAGA | TGGCAGGTGA |
| 5. | NL4-3:3405-3541 | + | 106 | 5.18e-7 | TATTATGACC | CATCAAAAGACTTAATAGCAGAAATA | CAGAA |
| 12. | NL4-3:7606-7743 | + | 27 | 6.57e-7 | CTGGAGGAGG | CGATATGAGGGACAATTGGAGAAGTG | AATTATATAA |
| 2. | NL4-3:1052-1188 | + | 33 | 1.50e-6 | GACACCAAGG | AAGCCTTAGATAAGATAGAGGAAGAG | CAAAACAAAA |
| 10. | NL4-3:6580-6717 | + | 57 | 1.99e-6 | ATGATACTAA | TACCAATAGTAGTAGCGGGAGAATGA | TAATGGAGAA |
| 14. | NL4-3:9221-9367 | + | 88 | 2.99e-6 | TGGAATGGAT | GACCCTGAGAGAGAAGTGTTAGAGTG | GAGGTTTGAC |

**C**

**SRSF motif**

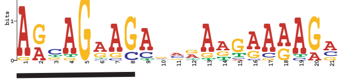

| | Name | Strand | Start | *p*-value | Sites | | |
|---|---|---|---|---|---|---|---|
| 12. | GX2005002:8041-8175 | + | 110 | 1.29e-8 | TCGCAGGACC | AGCAGGACAGGAATGAAAAGG | ATTT |
| 3. | GX2005002:2111-2268 | + | 101 | 3.75e-8 | GACTTACTGA | AGCAGGAGCAGAAAGACAAGG | AACATCAGCC |
| 4. | GX2005002:2580-2725 | + | 49 | 5.23e-8 | AGTGGCCATT | GACAGAAGAGAAAATAAAAGC | ATTAACAGAA |
| 10. | GX2005002:7086-7222 | + | 103 | 5.82e-8 | CAGGAGAAAT | AATAGGAGATATAAGAAAAGC | ATATTGCGAA |
| 15. | GX2005002:9242-9366 | + | 5 | 1.46e-7 | GTAG | AGGAGAACACCAAAGGAGAAA | ACAACTGCCT |
| 9. | GX2005002:6156-6292 | + | 57 | 2.15e-7 | TAAGAGAAAG | AGCAGAAGACAGTGGAAATGA | GAGTGAAGGA |
| 1. | GX2005002:719-856 | + | 66 | 2.59e-7 | AATTTTGACT | AGCGGAGGCTAGAAGGAGAGA | GATGGGTGCG |
| 11. | GX2005002:7624-7762 | + | 76 | 3.11e-7 | CACCAGGGCA | AAGAGAAGAGTGGTGGAGAGA | GAAAAAAGAG |
| 7. | GX2005002:4964-5097 | + | 41 | 3.11e-7 | GTGATATAAA | AGTAGTACCAAGAAGAAAAGC | AAAGATCATT |
| 8. | GX2005002:5505-5656 | + | 44 | 4.82e-7 | TTAAGAAATT | AACAGAAGATAGATGGAACAA | GCCCCAGAGG |
| 2. | GX2005002:1061-1154 | + | 68 | 5.26e-7 | GAGGGAGAAC | AAAAGAAGAGCCAGCAAAAGA | CACAG |
| 14. | GX2005002:8854-8991 | + | 7 | 7.96e-7 | TCCTCC | AGCAGCAGAAGGAGTAGGAGC | AGTATCTCAA |
| 6. | GX2005002:4063-4197 | + | 51 | 7.96e-7 | TCAACCAAAT | AATAGAGGAGCTAATAAAAAA | GGAAAAGGTC |
| 13. | GX2005002:8388-8470 | + | 19 | 2.47e-6 | TCGAAGAAGA | AGGTGGCGAGCAAGGCAGAGA | CAGATCCGTG |
| 5. | GX2005002:3440-3572 | + | 89 | 3.04e-6 | AAGACTTAGT | AGCAGAAGTACAGAAACAAGG | GCAGGACCAA |

### Supplementary Figure S10

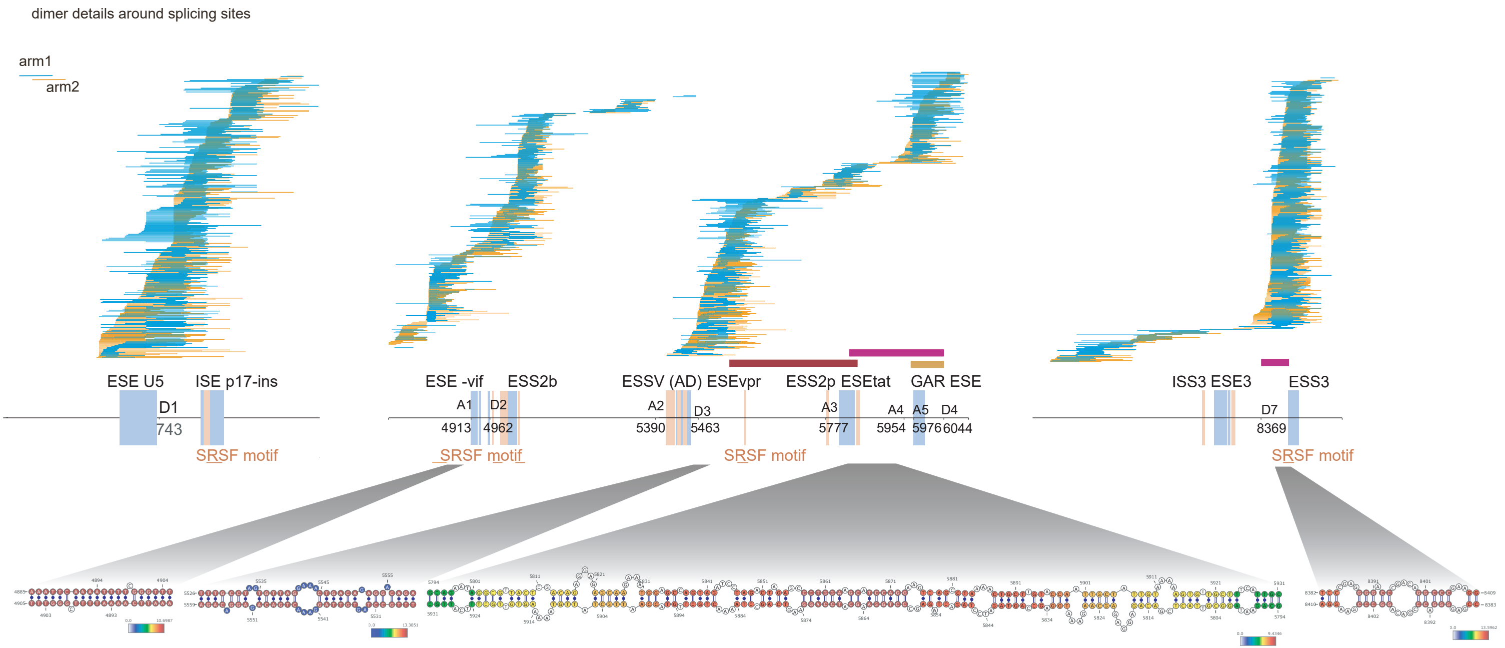

### Supplementary Figure S11

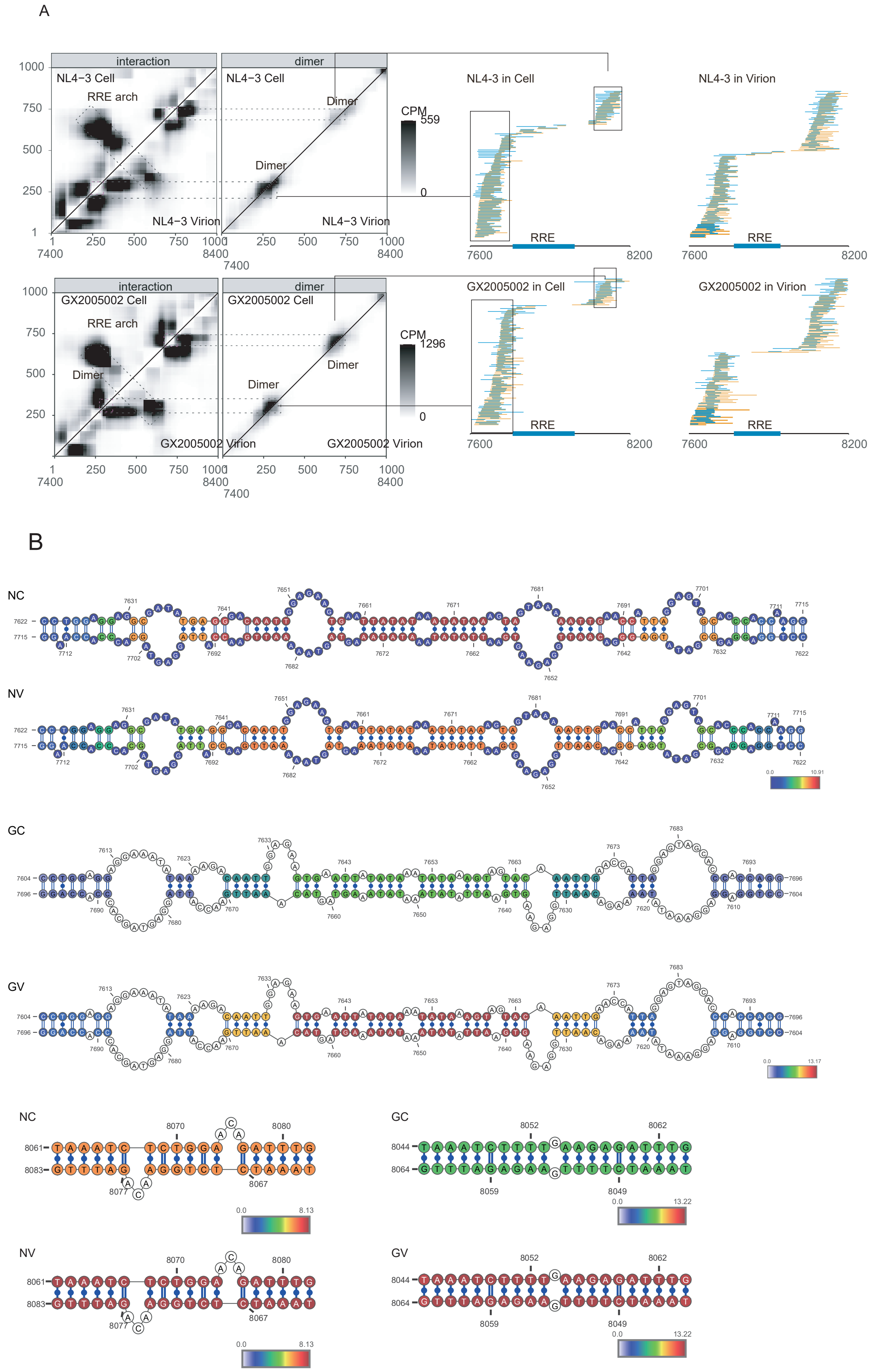

### Supplementary Figure S12

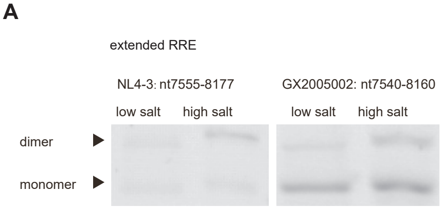


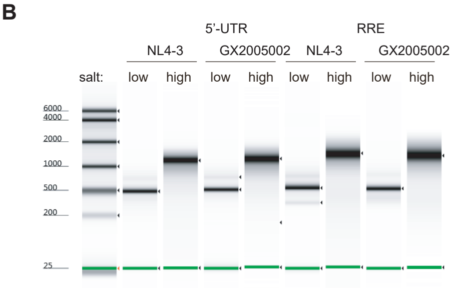

Supplementary Figure S13

#### NL4-3 RRE 378nt

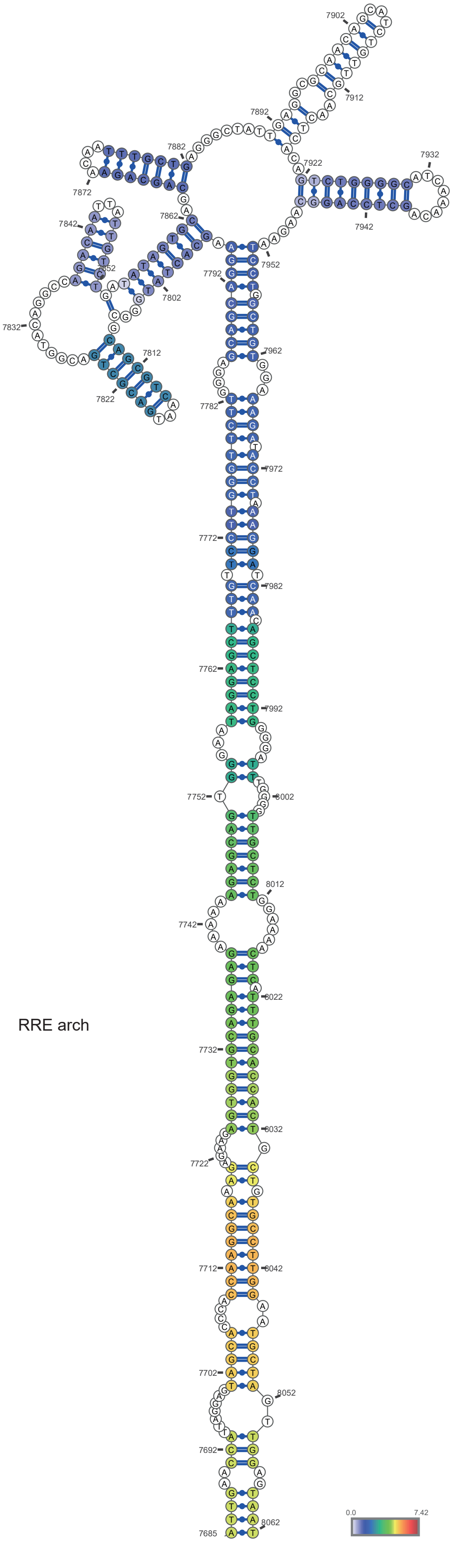


NL43: nt7685-8062 (E=-131.1 kcal/mol)

#### GX2005002 RRE 368nt

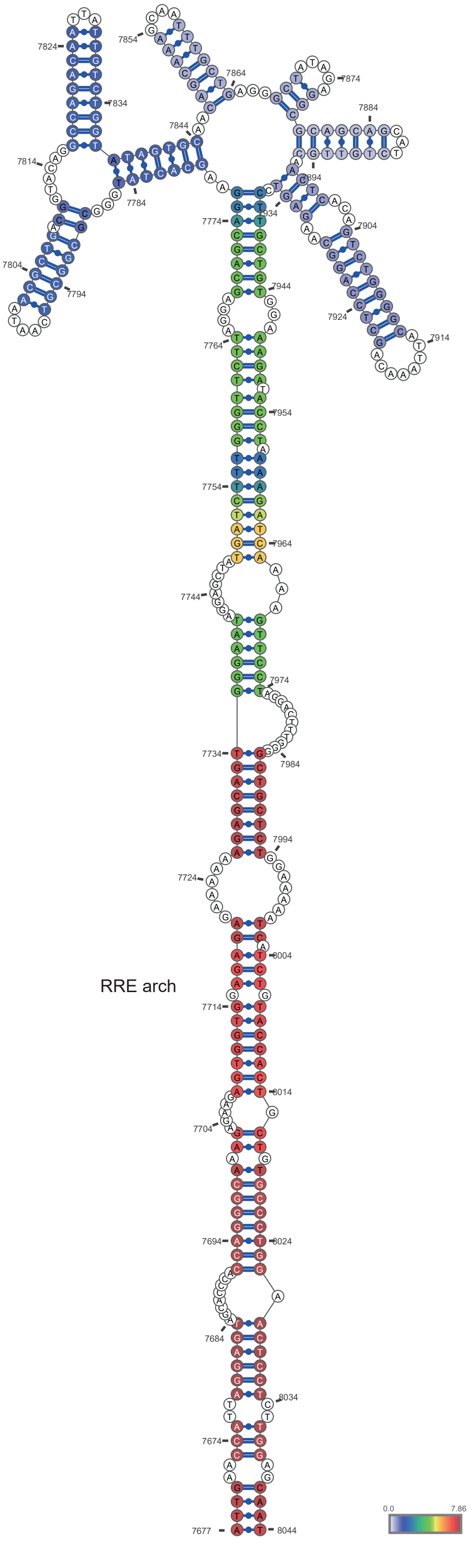


GX2005002: nt7677-8044   (E=-132.4 kcal/mol)

### Supplementary Figure S14

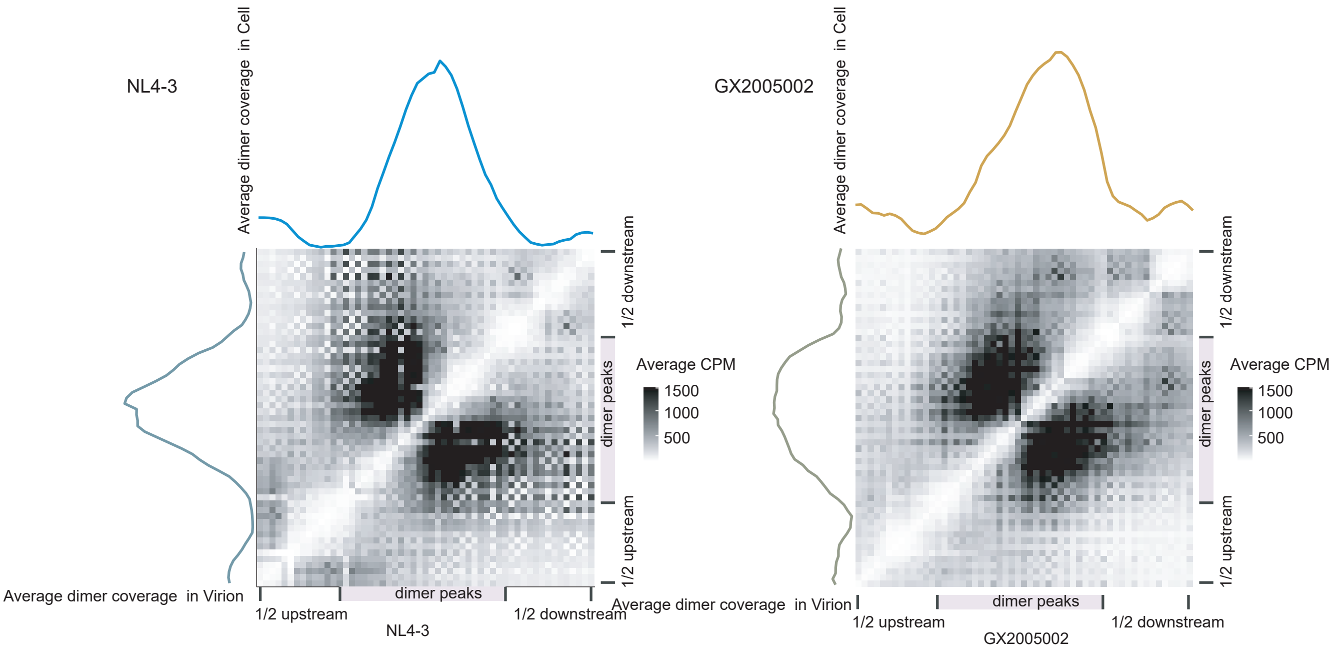

### Supplementary Figure S15

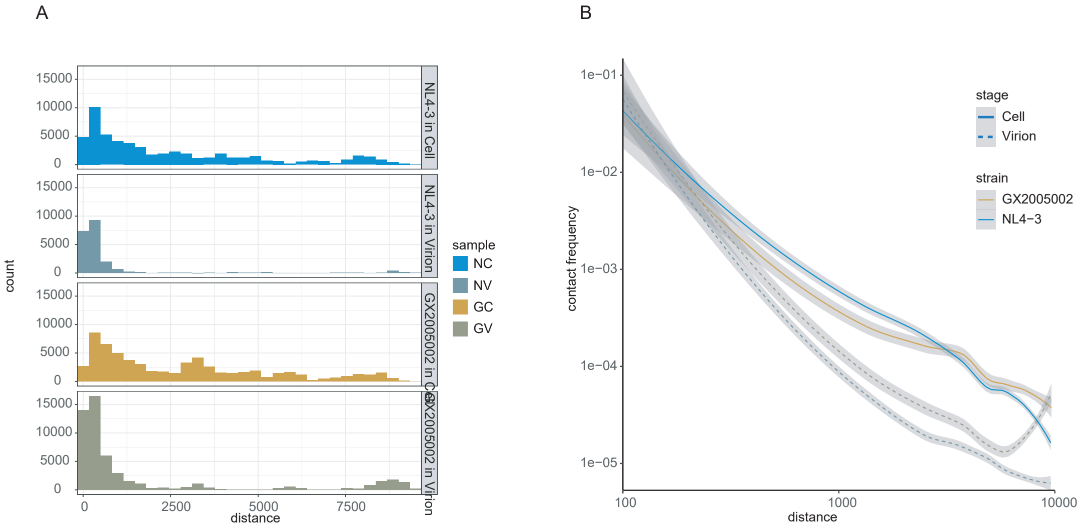

#### Supplementary Figure S16

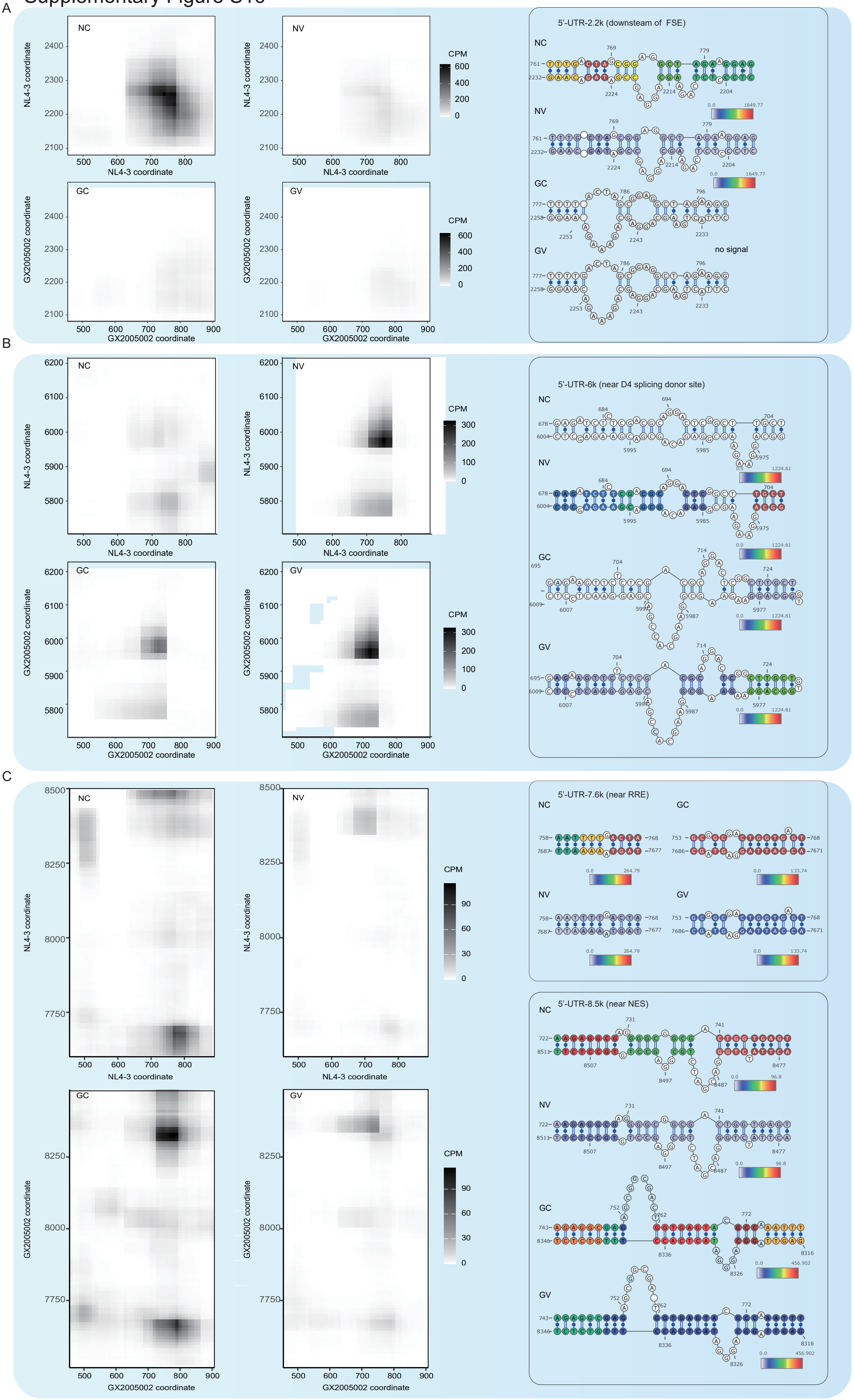
